# Somatostatin-expressing neurons in the ventral tegmental area innervate specific forebrain regions and are involved in the stress response

**DOI:** 10.1101/2023.01.06.522982

**Authors:** Elina Nagaeva, Annika Schäfer, Anni-Maija Linden, Lauri V. Elsilä, Maria Ryazantseva, Juzoh Umemori, Ksenia Egorova, Esa R. Korpi

## Abstract

Expanding knowledge about the cellular composition of subcortical brain regions demonstrates large heterogeneity and differences from the cortical architecture. Recently, we described three subtypes of somatostatin-expressing (Sst) neurons in the mouse ventral tegmental area (VTA) and showed their local inhibitory action on the neighbouring dopaminergic neurons (Nagaeva et al., 2020). Here, we report that VTA Sst neurons especially from the anterolateral part also project far outside the VTA and innervate several forebrain regions that are mainly involved in the regulation of emotional behaviour. Deletion of these VTA Sst neurons affected several behaviours and drug responses, such as home cage activity, sensitization of locomotor activity to morphine, fear conditioning responses, and reactions to inescapable stress of forced swimming, often in a sex-dependent manner. Together, these data demonstrate that VTA Sst neurons have selective projection targets, which are distinct from the main targets of VTA dopamine neurons and involved in the regulation of a variety of behaviours mostly associated with the stress response. This, makes Sst neurons a meaningful addition to the remote VTA circuit and stress-related neuronal network.

## Introduction

The ventral tegmental area (VTA) is a part of the midbrain, from which it sends neuronal projections to many brain structures. It is mainly recognized as the origin of two important dopaminergic pathways: the mesolimbic pathway to the ventral striatum and the mesocortical pathway to the prefrontal cortex, which control motivation and reward-related processes (Björklund and Dunnett, 2007). However, the VTA additionally contains two other major neuronal subtypes: neurons releasing glutamate (Glu) and γ-aminobutyric acid (GABA). In addition to participating in local circuits and controlling the neighbouring dopamine (DA) cells (Dobi et al., 2010; Hnasko et al., 2012; Tan et al., 2012), some of these neurons can project outside the VTA and contribute to larger brain circuits. It was shown that VTA GABA neurons can project to distant brain areas and modulate the activity of those areas separately from DA signalling, by having unique projection targets (Bouarab et al., 2019). For example, continuous activation of inhibitory projections from the VTA to the epithalamic lateral habenula promotes rewarding behaviours independently of DA neurons (Stamatakis et al., 2013). At the same time, activation of the rostral VTA GABA neurons projecting to the GABA neurons in the dorsal raphe disinhibits serotonin neurons and promotes aversion (Li et al., 2019). On the other hand, many VTA Glu projections travel in parallel with DA pathways (Cai and Tong, 2022), suggesting their sustaining role in DA signalling. It has also been shown that VTA Glu neurons projecting to the nucleus accumbens (NAc) could drive positive reinforcement and promote wakefulness independently from the DA release (Yu et al., 2019; Zell et al., 2020).

Recently, we described three VTA Sst+ neuron populations with heterogeneous molecular profiles (75% GABA, 18% Glu, and 5% GABA/Glu), location within the VTA and electrophysiological properties (Nagaeva et al., 2020). We also demonstrated that laterally located VTA Sst neurons were able to inhibit neighbouring DA cells via direct GABAergic transmission. In the present study, we demonstrate that anterolateral VTA Sst neurons project to distant forebrain regions and that deletion of these cells affects stress responses, often in a sex-dependent manner. Interestingly, our data suggest that the majority of the projecting Sst neurons was positive for *Vglut2* and *Th*.

## Results

### VTA Sst neurons innervate several forebrain regions

To find out whether Sst neurons project outside the VTA, we injected a Cre-dependent anterograde tracer unilaterally into the VTA of Sst-Cre mice and sectioned the whole brain three weeks later to locate the GFP signal of the tracer. We found VTA Sst projections in several forebrain regions. Then, we defined the regions, which consistently had the densest axonal arborisations in all brains studied, using hierarchical clustering and depicted the results as a heatmap (Fig. 1). VTA Sst neurons were found to have five main consistent projection targets: the ventral pallidum (VP), lateral hypothalamus (LH), the medial part of the central amygdala (CeM), anterolateral division of the bed nucleus of stria terminalis (alBNST), and paraventricular thalamic nucleus (PVT) (Fig. 1 and 2).

**Figure 1.**
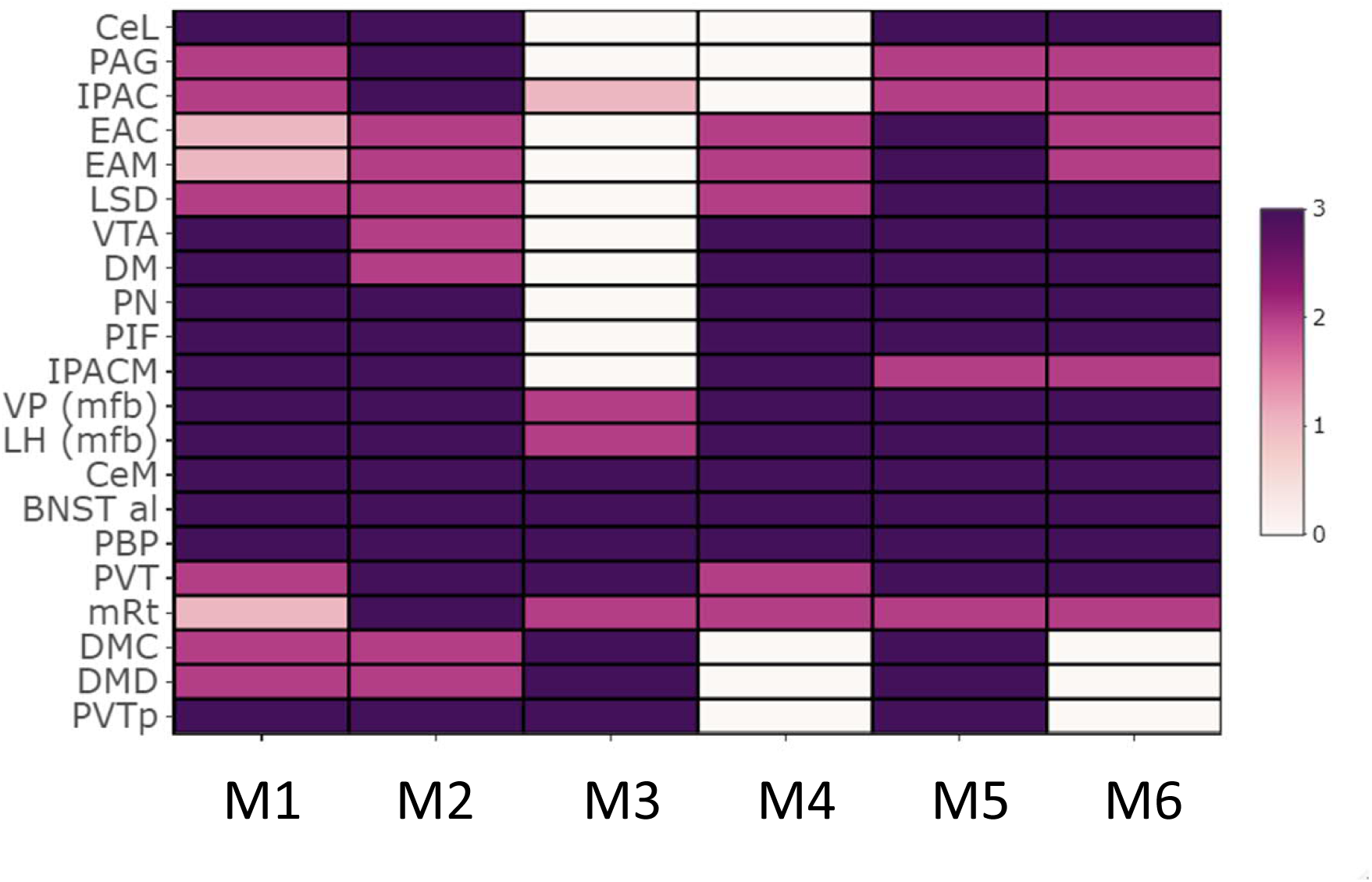
Heatmap of the VTA Sst neuron projections. The cluster of brain regions with the densest innervation is shown. Colours indicate the density of the fluorescent axons in the projection site from 0 (no projections) to 3 (highest density). The Y-axis depicts the names of the regions, X-axis shows the mouse number: M1 – mouse 1 and etc. The script and the source data to reproduce the clustering can be found here: https://github.com/eLinanin/Anterograde_heatmap.git ***BNST al*** – bed nucleus of the stria terminalis, antero-lateral part; ***CeL*** – central amygdala, lateral part; ***CeM*** – central amygdala, medial part; ***DM*** – dorsomedial hypothalamic nucleus; ***DMC*** – DM, compact part; ***DMD*** – DM, dorsal part; ***EAC*** – extended amygdala, central part; ***EAM*** – extended amygdala, medial part; ***IPAC*** – interstitial nucleus of the posterior limb of the anterior commissure; ***IPACM*** – IPAC, medial part; ***LH*** – lateral hypothalamus; ***LSD*** –lateral septal nucleus, dorsal part; ***mRT*** – mesencephalic reticular formation; ***PAG*** – periaqueductal gray; ***PBP*** – parabrachial pigmented nucleus of the VTA; ***PIF*** – parainterfascicular nucleus of the VTA; ***PN*** – paranigral nucleus of VTA; ***PVT*** – paraventricular nucleus of the thalamus; ***PVTp*** – PVT, posterior part; ***VP (mfb)*** – ventral pallidum (medial forebrain bundle); ***VTA*** – ventral tegmental area.

**Figure 2.**
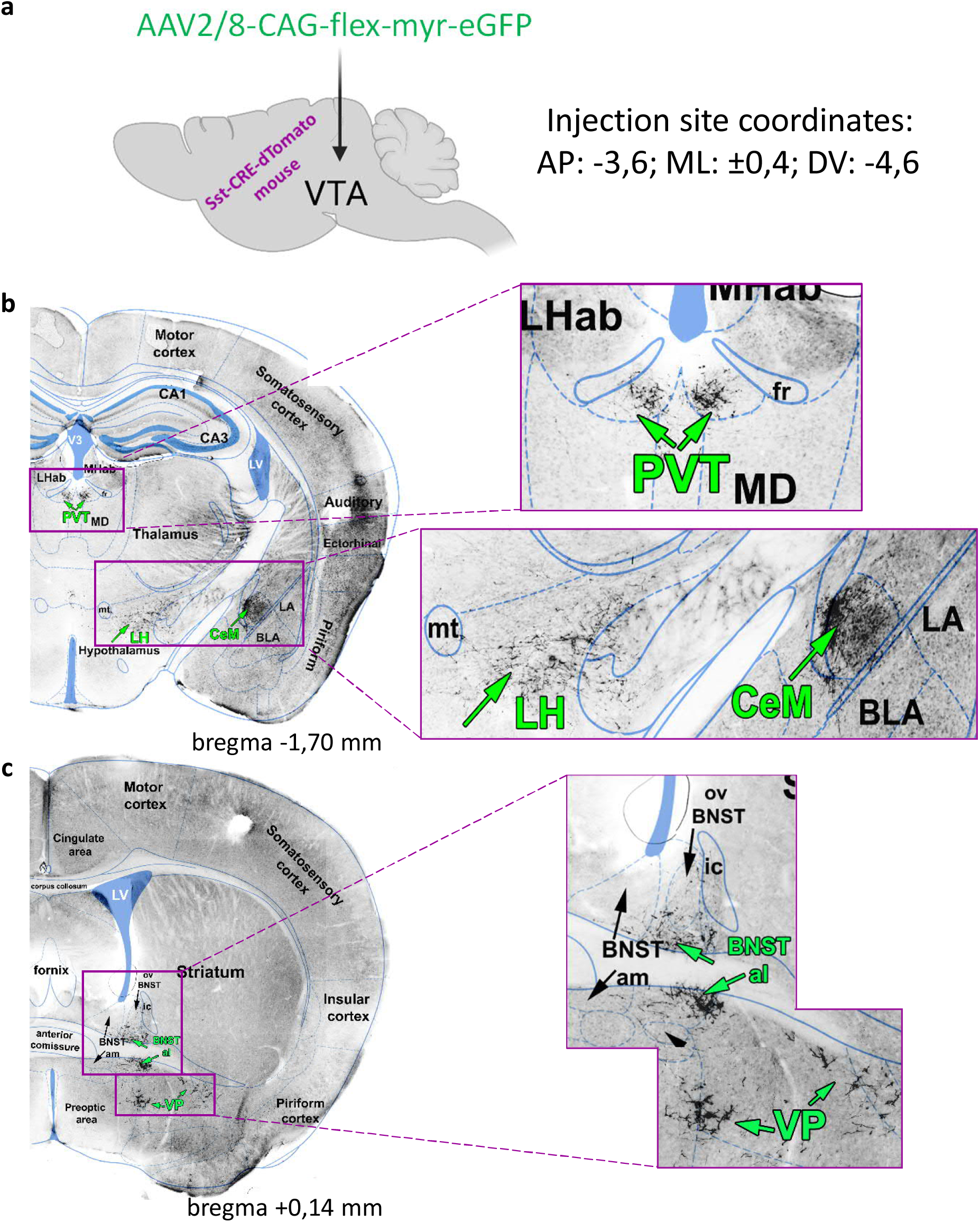
Anterograde tracing of the VTA Sst neurons. **a.** Sst-tdTomato mice received a unilateral intra-VTA injection of a Cre-dependent AAV tracer. The scheme shows the name of the viral tracer and injection coordinates. **b.** Examples of the VTA Sst+ projections found in the PVT, LH and CeM at the bregma level -1.70 mm. **c.** Examples of the VTA Sst+ projections found in the alBNST and VP at the bregma level +0.14 mm. **b-c** images are the black and white variants of the fluorescent GFP+ images of the coronal mouse brain sections.

Some of the projection targets, such as the LH and VP, are along the way of the medial forebrain bundle (mfb) – the massive neuronal tract connecting the midbrain with the forebrain. To confirm that VTA Sst axons innervate the targets mentioned and not only pass through, we injected a Cre-dependent retrograde tracer unilaterally into each of these targets (Fig. 3a). Indeed, injections into the CeM, LH, BNST and VP, produced GFP expression in the cell bodies of VTA Sst neurons located ipsilaterally to the injection site (Fig. S1-S4). However, unilateral injection of the retrotracer into the PVT produced GFP expression bilaterally in the VTA (Fig. 3b). Similarly, we detected traced axons in the left and right parts of the posterior PVT after unilateral VTA injection of the anterograde tracer (Fig. 2b). This might suggest that either individual VTA Sst neurons can send collaterals to the right and left parts of the PVT, or different Sst neurons from the same VTA side innervated the PVT bilaterally. Another explanation may lie in the non-bilateral anatomy of the PVT, which is a member of the midline thalamic nuclei family (Kirouac, 2015).

**Figure 3.**
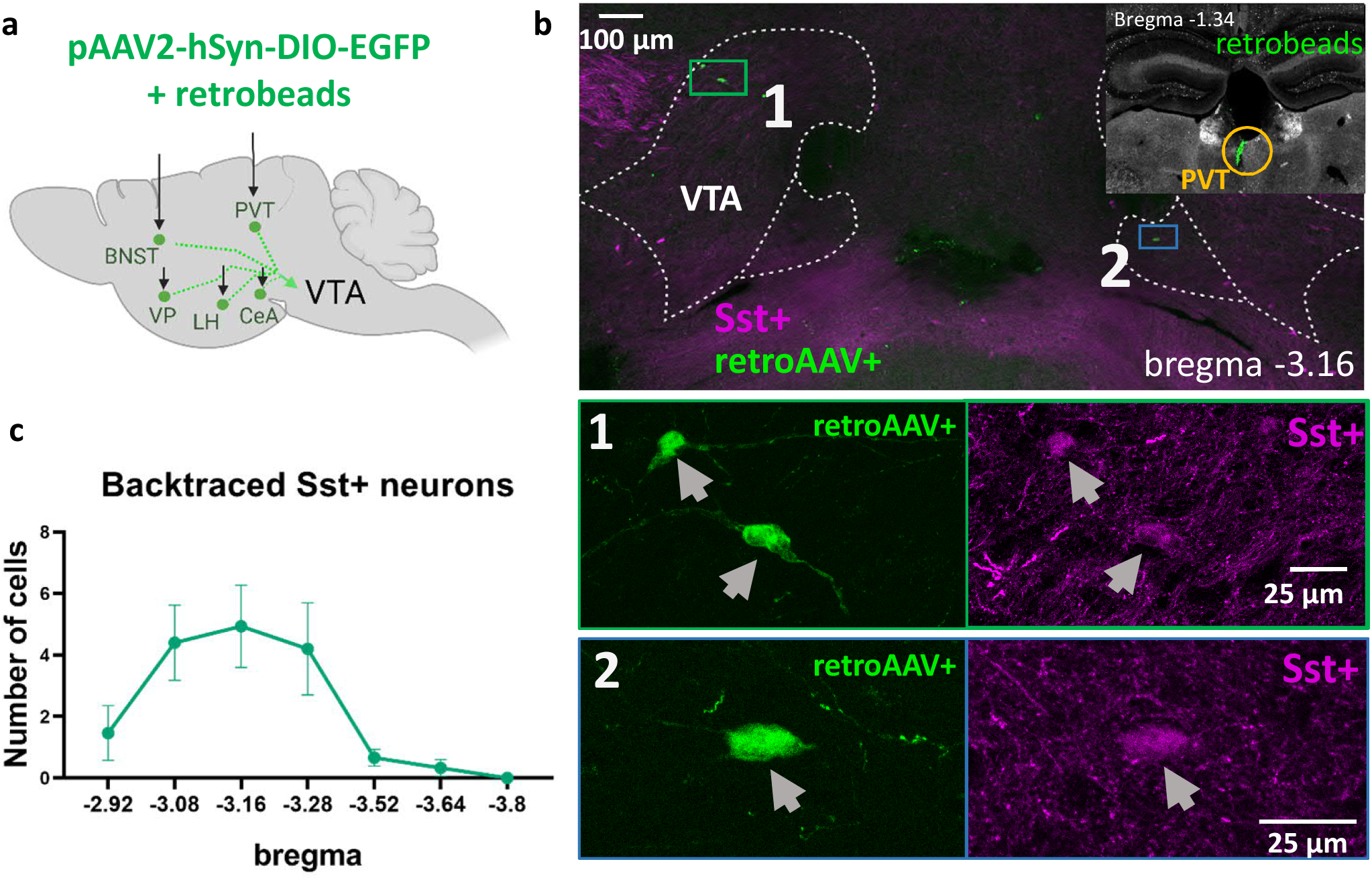
Retrograde tracing of the VTA projecting Sst neurons. **a.** Sst- tdTomato mice each received a unilateral injection of the mixture of the Cre-dependent retro-AAV virus and retrobeads in one of the five found VTA Sst projection targets (Fig.2). **b**. Examples of the backtraced neurons in the VTA at the bregma level -3.16 mm. The right upper corner shows retrobeads (green dots) in the injection site within the PVT (yellow circle). The green rectangle shows ipsilaterally traced neurons (1) and the blue rectangle shows a contralaterally traced neuron (2). Lower panels 1 and 2 are the magnified images of the green and blue rectangles. **c.** The graph shows the distribution of the backtraced VTA neurons at different bregma levels. The number of the backtraced Sst+ neurons from 5 different targets (LH, CeM, PVT, BNST and VP) were combined and are shown as average ± S.E.M. (n=15 mice).

### The majority of VTA Sst projecting neurons have electrophysiological ADP subtype and express *Th* and *Vglut2*

Retrograde tracing experiments also showed location specificity of VTA Sst projecting neurons in the VTA. Most (64/88 analyzed) of the backtraced neurons were found in the anterolateral part of the VTA at the bregma levels between −2.9 and −3.3 mm, with more than half of the cells found at the bregma levels -3.08 and -3.16 mm (Fig. 3c, 4b). We previously demonstrated that different electrophysiological subtypes of VTA Sst neurons have distinct locations, assigning anterolateral VTA neurons to either afterdepolarizing (ADP) or high-frequency firing (HFF) subtypes (Nagaeva et al., 2020). Therefore, we took an effort to define the electrophysiological subtype of the projecting neurons. To do that, we repeated the same procedure that was used for the backtracing experiments and performed patch-clamp recordings on GFP-positive VTA neurons. We also took advantage of the patch-clamp method to be combined with single-cell mRNA extraction (Sucher and Deitcher, 1995; Fuzik et al., 2016; Cadwell et al., 2017) and used the collected cell contents for further PCR analysis of the expression of main neurotransmitter markers.

**Figure 4.**
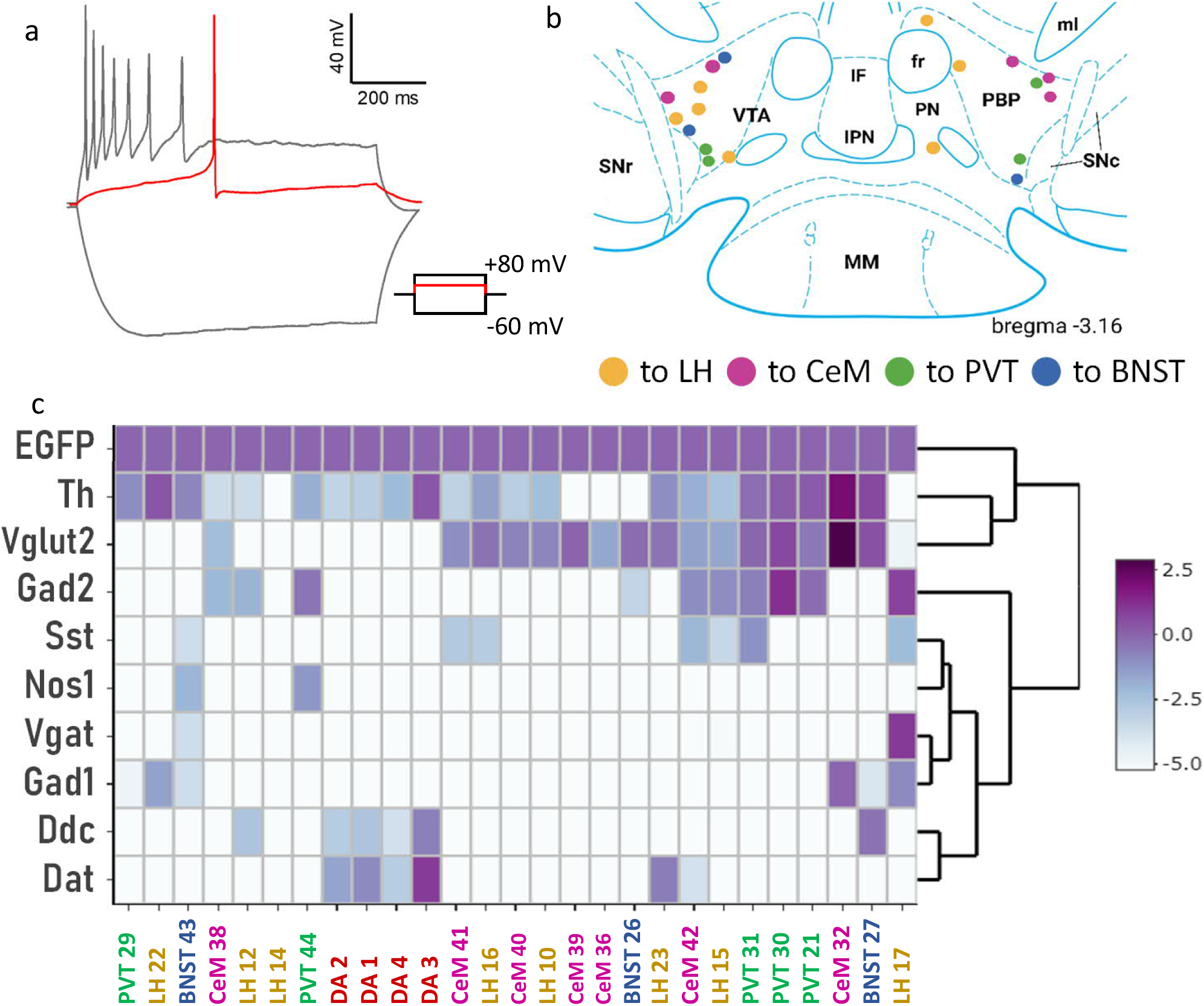
Electrophysiological and expression profiles of the VTA neurons projecting to the forebrain. **a.** A representative trace of the response of a VTA Sst projecting neuron to injected 800-ms current steps of −60, +8 (red line) and +80 mV. Adapting firing pattern at −80 mV and no delay before the firing resemble features of the ADP Sst subtype (see Nagaeva et al., 2020). **b.** Locations of the recorded VTA Sst neurons at the bregma level −3.16 mm. Their projection sites are colour-coded. **c.** The heatmap shows single-cell qPCR results for the main neurotransmitter markers of the backtraced VTA neurons. *EGFP* expression was used as a reference gene and has an expression value of 1. The majority of VTA Sst projecting neurons show expression of *Th* or *Vglut2*, or both, and lack of *Vgat* and *Dat*. Hierarchical clustering analysis did not indicate any neurotransmitter specificity depending on the projecting target. Cell names and colour codes on the X-axis represent the projection site of the cells. DA 1-4 are “control” dopamine cells from Th-EGFP mice and showed detectable expression levels of *Th*, *Dat* and *EGFP*. The script and the source data to reproduce the clustering at the Figure 3c can be found here: https://github.com/eLinanin/Fig4c_qPCR_heatmap.git

For the classification of electrophysiological subtypes of projecting Sst cells, we applied automatic firing pattern analysis (Nagaeva et al., 2021) and clustering algorithm (Nagaeva et al., 2020). We used our previously published electrophysiological dataset containing the firing patterns of 389 VTA Sst neurons as the reference dataset in the clustering procedure (for the full description of the method and reference dataset, see Nagaeva et al., 2020). The majority (67%) of the backtraced neurons were assigned to the ADP cluster (Fig. S5). Indeed, these neurons showed adapting firing rates at the saturated level of excitation, sag depolarization and small afterdepolarization in the first rheobase action potential (Fig. 4a), mimicking the ADP Sst subtype. Antero-lateral location of the recorded neurons (Fig. 4b) also supported their affiliation with the ADP subtype.

qPCR analysis of the recorded Sst neurons suggested that 78% (18/23 analyzed) of the neurons expressed tyrosine hydroxylase (*Th)* mRNA and 72% (16/23 analyzed) *Vglut2* mRNA with 13/23 cells having both *Th* and *Vglut2* transcripts (Fig. 4c). To address the question of possible contamination and to have a positive control for the DAergic nature, we also analyzed four DA neurons from the VTA of Th-EGFP mice using the same procedure (DA1-DA4 in Fig. 4c). These control DA cells expressed a combination of all classical dopaminergic markers, including *Th*, DOPA decarboxylase (*Ddc*) and dopamine reuptake transporter (*Dat*). None of the backtraced VTA Sst neurons was found to express the same full set of DA markers (Fig. 4c). Importantly, the control DA neurons did not express any other tested genes, suggesting a low level of contamination from surrounding cells during the cell collecting procedure. We were able to detect *Sst* mRNA in 30% of the recorded neurons, although, in our mouse model, GFP could be expressed in these neurons only via a Cre-dependent mechanism under the Sst promotor. This fact might be explained by the low level of Sst mRNA expression in Sst-Cre mice (Viollet et al., 2017) which however did not affect expression levels of other neuropeptides or Sst receptors in these animals. Only 2 out of 23 analyzed Sst projecting neurons showed expression of a combination of the two major GABAergic markers *Vgat* and *Gad1*. We did not observe any specificity in expression profiles depending on the projection target.

### Behavioural consequences of the deletion of VTA Sst neurons

We used “the loss of function” approach to elucidate the behavioural impact of VTA Sst neurons and selectively deleted them by injecting a Cre-dependent caspase-expressing virus bilaterally into the VTA area of adult Sst-Cre-tdTomato mice. After activation, this virus started the programmed cell death and eliminated Sst neurons exclusively in the injected region (Yu et al., 2019), resulting in “VTA**^Sst-^** mice”. To visualize successful deletion, the GFP-expressing Cre-dependent virus was injected either together with the caspase virus or alone into the control group (Fig. S6). We aimed to delete neurons preferably in the anterolateral VTA, where most of the projecting Sst neurons were found (see Fig. 3c).

For the proper planning of the behavioural experiments, we considered the previously established function of ADP Sst neurons as interneurons (Nagaeva et al., 2020) and the physiological function of their projection targets. Since, on one hand, VTA Sst ADP neurons can inhibit locally neighbouring DA neurons, we used several reward and motivation-related tasks to find out how the impairment of the local inhibitory circuitry affects these processes (Van Zessen et al., 2012; Corre et al., 2018). On the other hand, since they project to forebrain targets such as the PVT, CeM, LH, VP and BNST implicated in the processing of the aversive stimuli and defensive behaviours (Keifer et al., 2015; Lebow and Chen, 2016; Gao et al., 2020; Gomes-de-Souza et al., 2021), we used stress-, anxiety- and fear-related behavioural tests. It was also important to find out whether there were any sex differences in the affected behaviours, as some of VTA Sst projection targets, such as the PVT, BNST and LH, are known to be sexually dimorphic structures (Kim et al., 2017; López-Ferreras et al., 2017; Uchida et al., 2019).

### Increase in home-cage activity of VTA^Sst-^ female mice

Table 1 contains information on the behavioural experiments, the numbers of animals tested in the control and VTA**^Sst-^** groups, and whether or not we found statistically significant changes in behaviour. Of the 18 tests performed, 5 tests revealed differences between the VTA**^Sst-^** and control groups. Interestingly, some of the differences were sex-specific. The VTA**^Sst-^**females were more active in nose- poking to the water-containing doors in the corners during the first 14 h in the Intellicage environment than the control females (Fig. 5a). Their activity remained upregulated after 39-62 h of adaptation to the Intellicage, suggesting that differences were not due to exploration of the novel environment. There were no changes in this activity between the control and VTA**^Sst-^** male mice (Fig. 5b). It is important to note that the number of licks to water bottle tips behind the doors was similar in all mouse groups, indicating no increase in water consumption in the VTA**^Sst-^** females, or in the females overall compared to males (data not shown).

**Figure 5.**
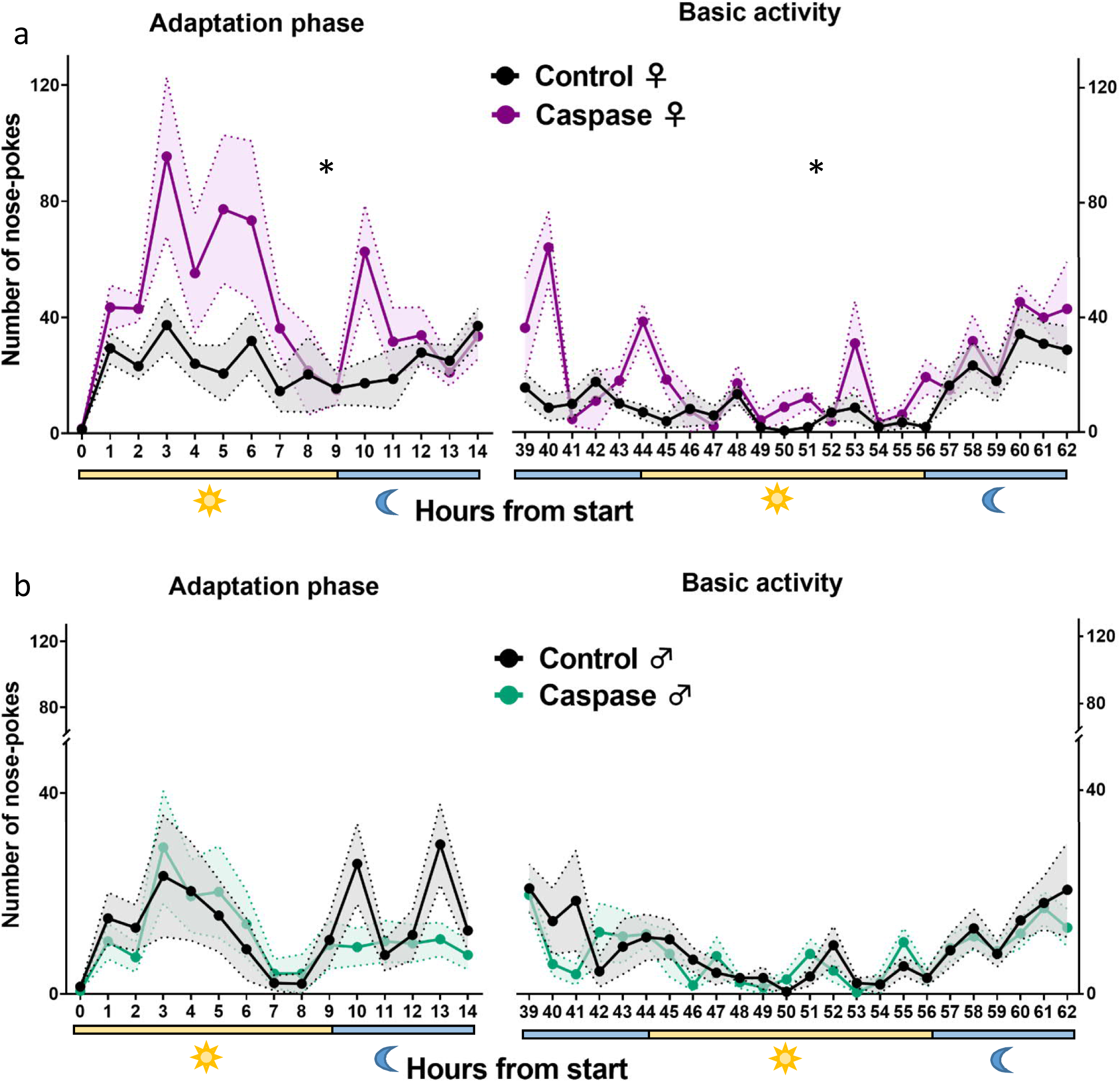
VTA^Sst^ -caspase female mice demonstrated an increased number of nose-pokes in the Intellicage system. Y-axis indicates the number of nose- pokes into the water-containing doors per h, X-axis shows the time after the beginning of the test. Yellow and blue bars beneath the graphs show light and dark phases, respectively. **a.** VTA^Sst^ –caspase females nose-poked more often in the Intellicage environment than the control females, suggesting higher activity during the adaptation (sex x treatment: F(1,22) = 7.085, p=0.014; ♀ *post-hoc* p=0.004) and after habituation to the new environment (sex x treatment: F(1,22) = 8.043, p=0.010; ♀ *post-hoc* p=0.002). Females were overall more active than males (sex: F(1,22) = 27.329, p<0.001). **b.** There was no difference between the male treatment groups. Data are shown as mean ± SEM,* p < 0.05 for the significance of the difference in overall activities (post-hoc test).

**Table 1.** Behavioural tests for VTA^Sst+^ and VTA^Sst-^ mice

| Name of the test | Environment | group size |  |  |  | Affected? | Statistical significance in |
| --- | --- | --- | --- | --- | --- | --- | --- |
|  |  | ♂ control | ♂ caspase | ♀ control | ♀ caspase |  |  |
| Running wheel | home cage | 9 | 6 | 8 | 5 | No | - |
| Open field | open arena | 10 | 9 | 11 | 8 | No | - |
| Light-dark box | test system | 10 | 9 | 11 | 8 | No | - |
| Elevated plus maze | test system | 10 | 9 | 11 | 8 | No | - |
| Novely suppressed feeding | open arena | 10 | 9 | 11 | 8 | No* | tendency to sex-treatment interaction; caspase males start eating faster than controls, and caspase females later than controls |
| Sucrose preference | home cage | 10 | 9 | 11 | 7 | No |  |
| Morphine-induced locomotor sensitization | open arena | 10 | 7 | 11 | 8 | Yes | expression of morphine sensitization, caspase mice display stronger morphine-induced locomotor sensitization |
| Forced swim test | test system | 10 | 9 | 11 | 8 | Yes | caspase animals struggle longer than controls |
| Saccharine preference | Intellicage | 7 | 7 | 7 | 5 | No |  |
| Home cage activity in group | Intellicage | 7 | 7 | 7 | 5 | Yes | females are more active in nose-poking to water-containing doors than males; caspase females are more active than control females. |
| Delay discounting | Intellicage | 7 | 7 | 7 | 5 | No* | Strong sex interaction; females are ready to wait longer for the saccharine than males. |
| Reward-related learning | Intellicage | 7 | 7 | 7 | 5 | No | - |
| Reward-related unlearning | Intellicage | 7 | 7 | 7 | 5 | No | - |
| Reward-related unlearning with punishment | Intellicage | 7 | 7 | 7 | 5 | No | - |
| Reward-related re-learning | Intellicage | 7 | 7 | 7 | 5 | No | - |
| Fear conditionning (FC) | test system | 6 | 7 | 5 | 5 | Yes | sex-treatment interaction; caspase males freeze less and caspase females freeze more than controls |
| FC context test | test system | 6 | 7 | 5 | 5 | Yes | sex-treatment interaction; caspase males freeze less and caspase females freeze more than controls |
| FC cue test | test system | 6 | 7 | 5 | 5 | No | - |

Similarly, we did not see any differences in the open-field and anxiety tests between the control and VTA**^Sst-^** mice (Fig. S8, Supplementary Table S3), suggesting that deletion of VTA Sst neurons influenced exclusively the home-cage activity in females and not the explorative activity or the level of anxiety.

### Fear conditioning is affected differently in VTA^Sst-^ males and females

Considering that VTA Sst neurons mostly project to the brain areas which control response and memory formation to aversive events (Keifer et al., 2015; Penzo et al., 2015; Goode and Maren, 2017; Concetti et al., 2020), we further assessed possible changes in threat processing. We used a Pavlovian fear conditioning paradigm (Maren, 2001), followed by contextual and cue-induced retrieval tests to assess differences in acquisition (acute response) and fear memory formation/expression. During the acquisition, mice were presented with three consecutive 30-s cue sounds (CS) co-terminated with 2-s foot shocks (0.6 mA) and separated with short breaks (Fig. 6a). Repeated measures two-way analysis of variance (ANOVA) for the per cent freezing and number of the freezing episodes detected a significant sex x treatment interaction (Supplementary Table S2, p=0.004 and p=0.035, respectively), indicating that the deletion of VTA Sst neurons influenced reaction to aversive foot-shock stimuli in a sex-dependent manner. A deeper analysis of the data showed that this difference came exclusively from the time points between the foot shocks when the sound was absent (breaks 1, 2 and 3; Fig. 6, Fig. S9b). Indeed, *post hoc* analysis confirmed that the VTA**^Sst-^**males froze less than the control males (p=0.036) during the breaks between foot shocks, whereas the VTA**^Sst-^** females froze more than the control females (p=0.038) (Fig. 6b). Interestingly, there were no differences between the VTA**^Sst+^** and VTA**^Sst-^** groups when the cue sound was on (Fig. S9b).

**Figure 6.**
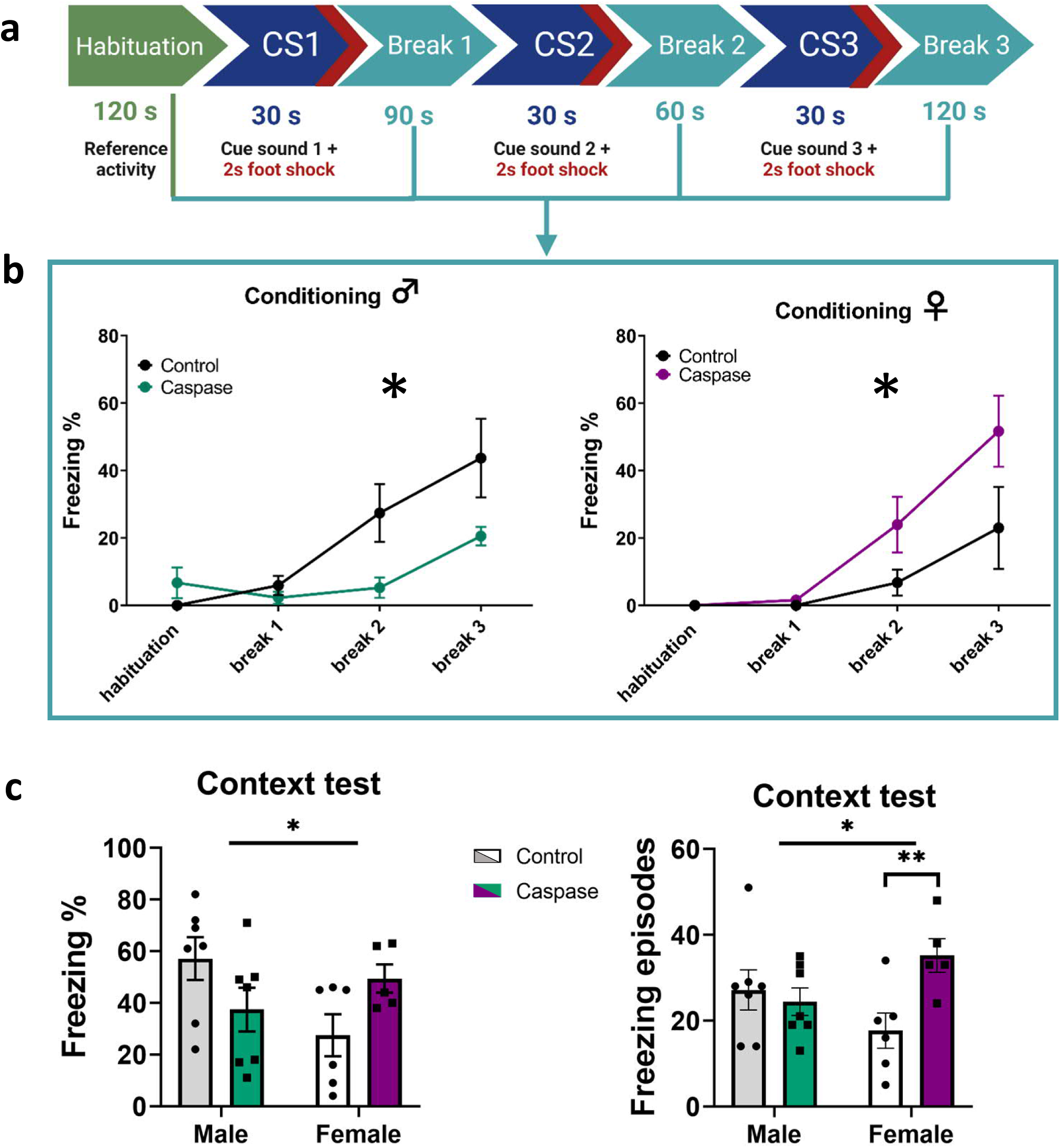
Deletion of VTA Sst neurons affected fear conditioning in a sex-dependent manner. **a.** Fear conditioning protocol during the acquisition phase (see methods). **b.** Graphs show per cent freezing (freeze time/total time) during breaks following the conditioning episodes (sex x treatment: F(1,21)= 9.971, p=0.005). The VTA**^Sst^**-caspase males froze less during the breaks between foot shocks, whereas the VTA**^Sst^**-caspase females, on contrary, froze more (*post hoc* males p=0.036, females p=0.038). **c.** A similar sex x treatment interaction (F(1,21)= 6.602, p=0.018 for freezing %; F(1,21)= 6.102, p=0.022 for freezing episodes) was observed in the context-associated fear memory retrieval test 9 days after the conditioning. The right graph shows that the VTA**^Sst^**-caspase females froze more often, but, as seen in the left graph, not significantly longer (freezing %) than the control females (freezing episodes *post hoc* females p=0.01). Data are shown as mean ± SEM. *p < 0.05, **p<0.01.

Further, we tested context-induced fear memory retrieval by placing the mice for 5 min in the same chamber, where they had received foot shocks 9 days earlier. Again, there was a similar sex x treatment interaction (p=0.018) in the per cent freezing and freezing episodes (p=0.022; Fig. 6c) as we saw during the conditioning. Although *post hoc* tests did not show significant differences in the per cent freezing time between the treatment groups within sexes, the VTA**^Sst-^** females showed more freezing episodes (p=0.01), meaning they froze more often but not significantly longer than the control females (Fig. 6c). Cue-induced retrieval or extinction after repeated cue presentations were not significantly affected by the deletion of VTA Sst neurons, although the results showed similar sex-dependent trends as during the conditioning and context testing (Fig. S9c).

### Deletion of VTA Sst neurons delayed the onset of immobility in the forced swim test

One of the non-sex-dependent changes in the behavioural performance of the mice lacking VTA Sst neurons was a delayed latency to the first immobility event in the forced-swim test (FST) (Fig. 7). Results showed that the VTA**^Sst-^** mice struggled longer than the control group (p=0.031) and tended to spend less time immobile during the first 4 min of the test. As the tested animals were not exposed to any chronic stressor before the FST, the observed behavioural alteration might predominantly be related to a reaction to the acute unpredictable stressor.

**Figure 7.**
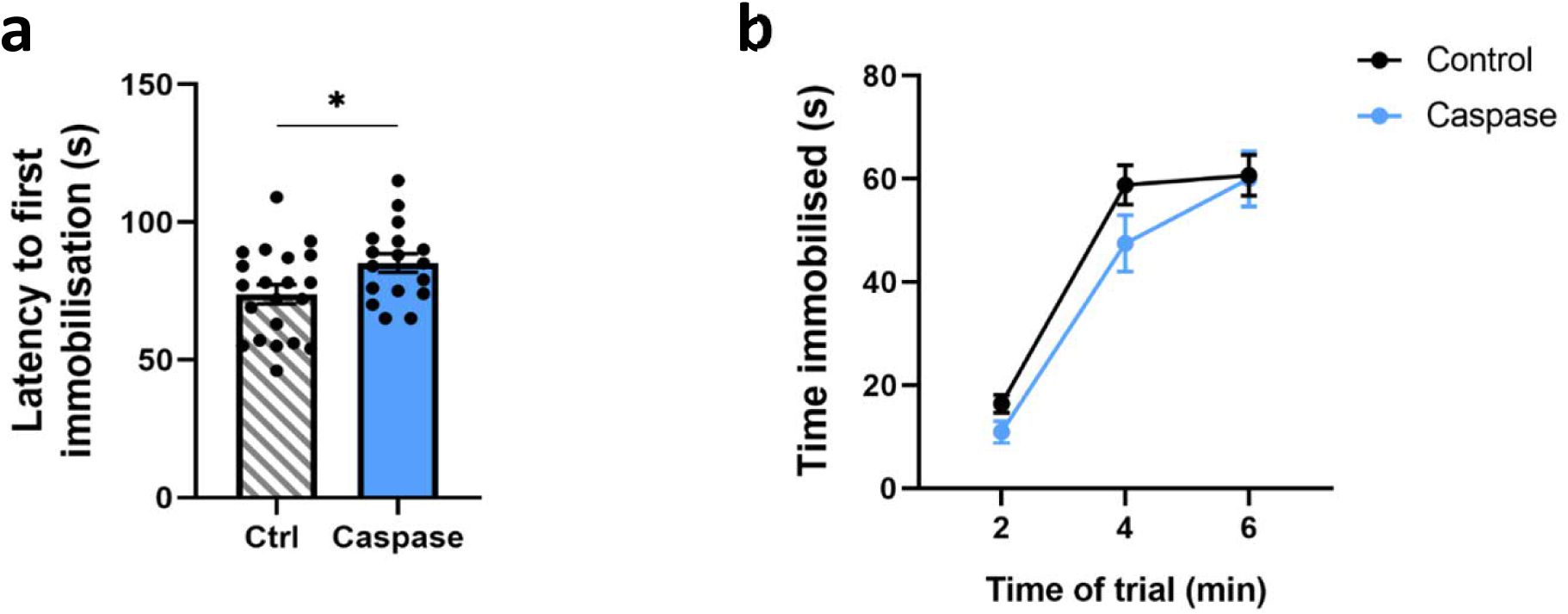
Delayed onset of immobility in the forced swim test in the VTA^Sst^-caspase mice. **a.** VTA**^Sst^**-caspase mice showed a longer latency to the first immobilization event (F(1,34) = 5.08, p = 0.031), **b.** and a tendency for a shortened duration of immobilization (time x treatment: F(2,68) = 1.938, p = 0.16) especially in the first four min. Data are shown as mean ± SEM, and dots in panel **a** show individual data points. *p < 0.05.

### Deletion of VTA Sst neurons affected morphine sensitization, but not natural reward-related processing

Taking into account the previously shown ability of VTA Sst neurons to inhibit neighbouring DA cells (Nagaeva et al., 2020), it was important to find out whether drug and natural reward processing would be affected in VTA**^Sst-^** animals. We chose morphine as the experimental substance since its rewarding potential is well-known in rodents (Martin et al., 1963; Kuzmin et al., 1996), and its mechanism of action includes inhibition of VTA GABA cells resulting in excessive DA neuron firing by disinhibition (Johnson and North, 1992). There were no differences in the locomotor response to a single-dose morphine administration (20 mg/kg, i.p.) between the treatment groups (Fig. 8a), indicating that acute reaction to morphine was not affected in VTA**^Sst-^** mice. However, we detected a significant enhancement in sensitization to locomotor activation by the second morphine challenge in the VTA**^Sst-^**mice as compared to the control mice, 7 days after the first morphine injection (Fig. 8b).

**Figure 8.**
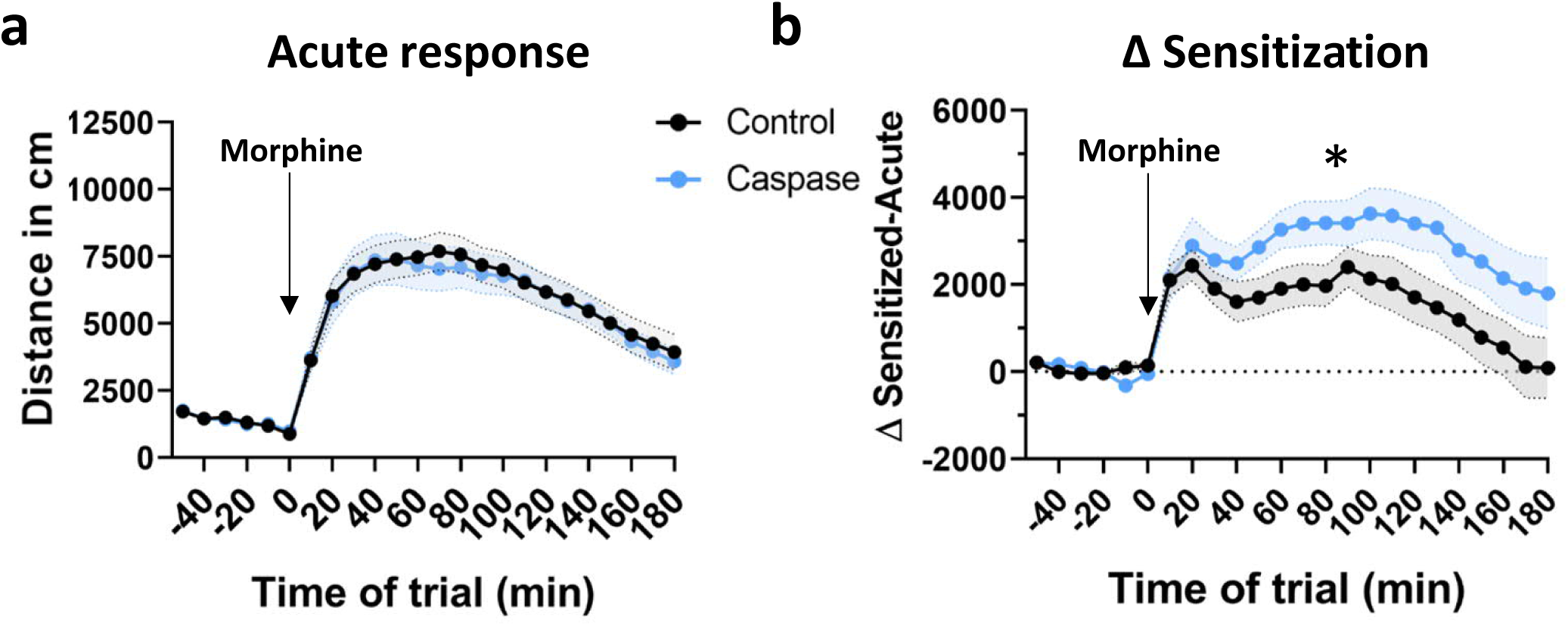
Increased morphine-induced locomotor sensitization in VTA^Sst-^ mice. Arrows indicate the time of morphine administration (20 mg/kg, i.p.) **a.** Acute treatment with morphine-induced similar hyperlocomotor response in both treatment groups (treatment: F(1,32) = 0.034, p = 0.854) **b.** Seven days after the first morphine injection, the response to the morphine challenge was enhanced and prolonged in the VTA**^Sst^**- caspase mice as compared to controls [Treatment: F(1,32) = 12.014, p = 0.002; Time x treatment: F(23,735) = 2.915, p = 0.024]. Data are shown across 10 min time bins as means ± SEM. *p < 0.05.

As the natural reward to be tested, we chose sweetened drinking water but did not find any significant differences between the treatment groups in sucrose or saccharine consumption or preference over plain water (Fig. S10). We also designed two reward-based learning tasks for the Intellicage system aiming to assess alterations in prediction error processing (Schultz et al., 1997) and in punishment- resistant reward preference. For these tasks, the mice learned first to nose-poke in assigned corners to receive access to 0.3% saccharine in water. Each nose-poke to the saccharine door activated light informing that the saccharine will be available in 2 s. On the third day, when the task performance was stable, we either emptied the saccharine bottles or introduced 0.2-bar air puffs with a 25% probability at the end of the licking session (Radwanska and Kaczmarek, 2012). It is important to note that the two experimental tasks took place consecutively in the Intellicage allowing assessment of the re-learning rates after failed reward accesses.

The “prediction error” probing showed significant sex differences (p<0.001) and sex x treatment interaction (p=0.043) between the control and VTA**^Sst-^** groups (data not shown). However, further *post hoc* analysis separately for males and females did not show any significant differences between the treatments, but showed a tendency for the VTA**^Sst-^** females to re-learn slower not to nose-poke anymore in an emptied saccharine bottle (p=0.064). For the next “punishment- resistant reward preference test”, saccharine was re-introduced in new corners after being unavailable for 2 days and the mice had to re-learn the rules. Although there was a tendency of the VTA**^Sst-^** male mice to be less active in nose-poking to saccharine corners in all phases of the re-learning/avoidance test (Fig. S11), we could not detect any statistically significant differences between the groups (see Supplementary Table S1). Similarly, we did not detect any differences in reactions to air puffs.

### Delay discounting test revealed a sex-dependent, but not VTA Sst neuron- dependent difference

Even though we did not find any differences in the preference for saccharine or sucrose of various concentrations between the VTA**^Sst-^** and control groups (Fig. S10), we observed an interesting sex-dependent behaviour in the delay discounting (DD) task (Mitchell, 2014) in Intellicage system. The delay discounting test, where mice learn to wait for a reward for increasing periods of time, showed a sex difference (p=0.0239) with significant dependence on the duration of the delay (sex x delay interaction p<0.001; Supplementary Table S1). As shown in Fig. 9, for both male groups the longer delay to the saccharine delivery drastically decreased the number of licks to the saccharine bottle starting from the 4-s delay and dropped almost to nonexistent at the 5-s delay, while females were still willing to wait and lick. The delay-discounting test is usually interpreted as a measure of impulsivity, making male mice in our experiment more impulsive or less patient. However, we did not see any differences between the treatment groups within the sexes, suggesting that the deletion of VTA Sst neurons did not affect impulsivity or readiness to wait for the reward.

**Figure 9.**
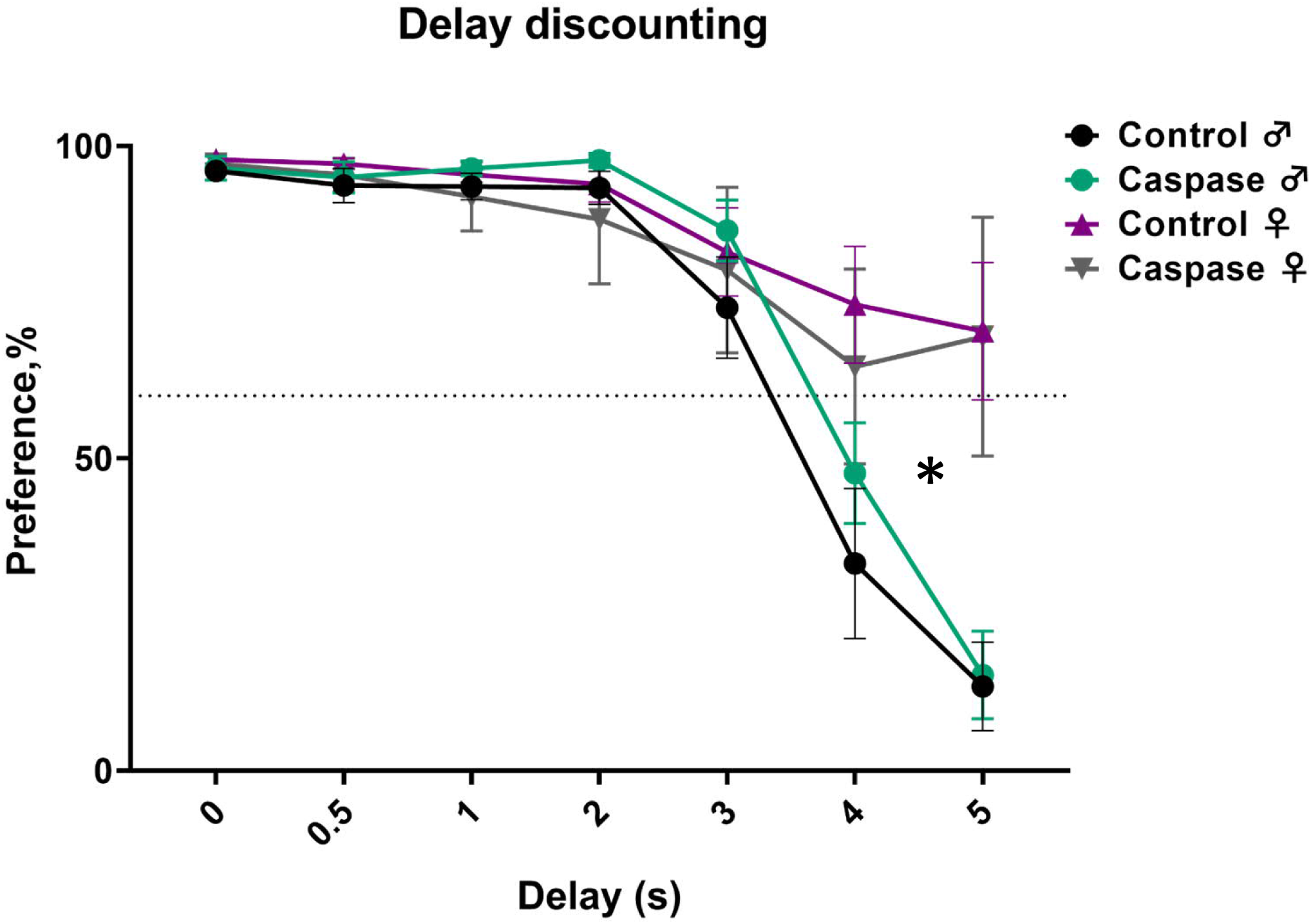
Deletion of the VTA Sst neurons did not influence the impulsivity or readiness to wait for the saccharine reward. Y-axis indicates preference in % for 0.3% saccharine over water defined as a lick number to the saccharine bottle/total number of licks. The X-axis indicates the duration of the delay before the saccharine door opened after a mouse entered the corner. * indicates the significance of the differences between the sexes (F(1,22)=5.829, p=0.024) and of the sex x delay interaction (F(6,132)=17.09, p<0.0001). Data are shown as mean ± SEM.

## Discussion

In the present study, we demonstrated that Sst neurons similarly to other neurons in the VTA can project outside the midbrain. In addition to their local inhibitory activity onto neighbouring DA cells (Nagaeva et al., 2020), VTA Sst neurons consistently project to five forebrain targets: alBNST, CeM, PVT, LH and VP. These projecting Sst neurons have a specific location in the anterolateral part of the VTA, where most of the Sst neurons of the afterdepolarizing (ADP) subtype are located (Nagaeva et al., 2020). Here, we also confirmed the ADP elctrophysiological subtype for the backtraced Sst projection neurons by patch-clamp experiments. Our qPCR single-cell experiments suggested that many of projecting Sst neurons expressed *Vglut2* or *Th*, or both of them. The existence of double positive Th+/Vglut2+ neurons is well established in the VTA of mice and rats by several studies (Li et al., 2013; Yamaguchi et al., 2015), reporting also that about 50% of these neurons lack mRNAs for the main DA markers (confirmed by our qPCR results). The existence of a sizable *Vglut2+* Sst population was reported in our previous work (Nagaeva et al., 2020) by *in situ* hybridization (RNAscope) and patch-seq (Cadwell et al., 2015; Fuzik et al., 2016) approaches. This allows to suggest that projecting VTA Sst neurons may represent a subset of *Vglut2+* Sst neurons described previously.

Taking into account the mixed neurotransmitter nature of VTA Sst neurons and the fact that *Sst* can be expressed also in excitatory neurons in the subcortical areas (Phillips et al., 2022; Sun et al., 2022), it is possible that at least some of the targets found belong to the known VTA Glu neuron projections. For instance, some of VTA Vglut2+ neurons have been shown to activate and inhibit VP neurons with the excitatory action being more pronounced (Hnasko et al., 2012; Yoo et al., 2016). It is plausible that the Sst projections described here represent a part of these Glu projections.

On the other hand, optogenetic activation of the VTA neurons in Gad2-Cre mouse line showed a functional inhibitory connection of these neurons with neurons in the CeM (Zhou et al., 2019), suggesting that VTA Sst+ projections might constitute at least a part of that VTA-CeM pathway. The fact that ∼40% of the backtraced VTA Sst neurons showed expression of *Gad2* supports this possibility. Furthermore, projections of the VTA GABA and Glu neurons to the LH have been shown to regulate wakefulness and sleeping behaviour (Yu et al., 2019). Taking into account that the Sst neurons projecting to the LH expressed *Vglut2* and sometimes *Gad1*/*Vgat*, it is plausible that in the work of Yu et al. a sizable population of the Sst neurons described here was responsible for the regulation. Indeed, a more recent work from the same laboratory showed that VTA Sst GABAergic projections to the LH regulate restorative sleep after stress (Yu et al., 2022).

Behavioural assessment of the deletion of VTA Sst neurons revealed four major consequences: increased home-cage activity, altered response in the fear-conditioning, enhanced locomotor sensitization to the second dose of morphine and prolonged struggle during the inescapable stress of forced swimming. The first two changes showed a significant sexual dimorphism.

Regarding the home-cage activity, it is well known that females are more active than males in the Intellicage environment where they are group-housed (Pernold et al., 2019, 2021), as we also observed (Supplementary Table S1). Interestingly, the deletion of VTA Sst neurons resulted in a further increase of the home-cage activity exclusively in the female mice (Fig. 5). The activity change was not associated with the novelty of the environment: it was consistent throughout the experiment and after three days of adaptation. Circadian patterns in single-housed animals determined during 3-day access to free-running wheels failed to differ between VTA**^Sst-^** and control animals (Fig. S7). In addition, there was no change in locomotor activity, as the distance travelled in the open field was similar between the two treatment groups. Our data suggest that the deletion of VTA Sst neurons resulted in a specific increase in home-cage activity and only in group-housed female mice. The underlying mechanism of this distinct change needs further investigation.

Although several VTA Sst neuron projection targets, like the BNST (Lebow and Chen, 2016), CeM (Shackman and Fox, 2016) and PVT (Kirouac, 2021), are involved in the brain anxiety network, we did not see any significant differences in anxiety- like behaviours in VTA**^Sst-^**mice. These behaviours were measured in light-dark box, elevated plus maze, novelty-suppressed feeding tests (Fig. S8) or reaction to the air-puffs (Fig. S11), and the results suggest that VTA Sst neurons are not involved in innate threat and avoidance of elevated or brightly lit areas. No change in the elevated plus maze and open-field tests in VTA**^Sst-^** mice was also reported by another group (Yu et al., 2022), confirming our findings. On the contrary, our data indicate that these neurons could regulate the responses to inescapable acute stressors, such as freezing after unexpected foot shock (fear-conditioning acquisition phase) or struggling when dipped into water (forced swim test).

Significant, but opposite changes in the reaction to the foot shock in males vs. females (Fig. 6b) was another interesting effect of VTA Sst neuron deletion. It is important to note, that our data do not exclude the possibility that the nociceptive reaction itself was altered in the VTA**^Sst-^** animals explaining why VTA**^Sst-^**males froze less and VTA**^Sst-^** females froze more than controls in response to the foot shock. Interestingly, a recent study described a pathway from the laterodorsal tegmentum (LDTg) via the VTA to the basolateral amygdala, inhibition of which reduced unconditioned freezing response to foot shocks in male mice, similar to what we observed in the VTA**^Sst-^** males (Broussot et al., 2022). Unfortunately, results on females were not reported in that study. Our data emphasize the importance of conducting experiments on both sexes, especially in case of reactions to aversive-stimuli (see also Cover et al., 2014). Fear-conditioning test highlighted an interesting feature of VTA Sst neuron function: these neurons were important for the contextual fear memory formation/retrieval, but not for the sound cue-associated responses, likely relating to differential processing of these two modalities of conditioned stimuli.

The fact that many brain regions receiving VTA Sst neuron projections belong to the extended amygdala suggests the involvement of these neurons in the action of addictive substances, such as opioids and alcohol (Koob et al., 2014a, 2014b). We did not see any differences in locomotor responses to acute morphine administration (Fig. 8a), indicating that the stimulating effect of an opioid was not affected by the deletion of VTA Sst neurons and that opioid-induced disinhibition of DA neurons (Johnson and North, 1992) was probably not mediated by the VTA Sst neurons. This is consistent with the data on rats, where morphine actions on VTA DA neurons could be prevented by silencing GABA neurons in the rostromedial tegmental nucleus (RMtg, also called the tail of VTA) (Jalabert et al., 2011), which neurons might be more meaningful for the acute disinhibitory action of opioids on VTA DA neurons. More importantly, we did observe that after the second morphine administration 7 days later, the VTA^Sst-^ mice of both sexes demonstrated robustly increased locomotor sensitization, suggesting that VTA Sst neurons normally limit the sensitization to opioids, but not acute effects of opioids. This might be linked to the morphine-induced adaptation of GABA_A_ receptor-mediated transmission onto VTA DA neurons (Nugent et al., 2007), which would be missing from intra-VTA synapses of Sst neurons in caspase-treated VTA^Sst-^ mice.

Interestingly, previous studies have shown that stressful events, including those of emotional nature, often result in a similar increase in opioid sensitization leading to higher morphine preference (Kalivas et al., 1986; Leyton and Stewart, 1990; Shaham et al., 1992; Kuzmin et al., 1996). Since it has been shown that the deletion of VTA Sst neurons disrupts normal restorative sleep after social defeat stress (Yu et al., 2022), we can speculate that in both cases VTA Sst neurons are involved in adaptive changes needed to return the VTA circuit to the baseline state. The fact that VTA Sst neurons appear to have protective properties in social stress and morphine sensitization asks for further research on their potential in preventing the development of opioid addiction.

In summary, our study has demonstrated that in addition to their role as local interneurons, VTA Sst cells can project outside the hosting region and innervate several forebrain areas. The VTA connection with alBNST and PVT through Sst+ neurons was shown here for the first time. Our behavioural experiments suggest that VTA Sst neurons are involved in stress-related reactions, which together with the established location of the projecting VTA Sst neurons, provides a solid background for future investigation of their remote circuitry function. Further research using opto- or chemogenetic methods will help to confirm the transmitter identities of the separate VTA Sst-expressing neuronal projections and uncover how their inhibitory or excitatory action on postsynaptic partners affect animal behaviour.

## Materials & Methods

### Animals

All experimental procedures were performed on male and female mice of heterozygous Sst-IRES-Cre (Sst^tm2.1(cre)Zjh^/J) genotype resulted from cross-breeding of Sst-IRES-Cre (Sst^tm2.1(cre)Zjh^/J) strain with Ai14 tdTomato reporter strain (B6.Cg- Gt(ROSA)26Sor^tm14(CAG-tdTomato)Hze^/J). Only the “backtracing-qPCR” part was performed in homozygous Sst-IRES-Cre mice to prevent leakage of tdTomato signal into the GFP-channel complicating the detection of the backtraced SSt eGFP-positive neurons Two female Th-eGFP mice (Tg(Th-EGFP)DJ76Gsat) of the age P140 were used for the qPCR experiment on the dopamine neurons. Animals were group-housed in individually ventilated cages (IVC)-cages (Tecniplast Green Line GM500) under 12:12 h light/dark cycle (lights on 6 am – 6 pm) with *ad libitum* access to food (Global Diet 2916C, pellet 12 mm, Envigo) and water, unless otherwise indicated in the corresponding method section. Cages were equipped with bedding (500 ml aspen chips 5 x 5 x 1 mm, 4HP, Tapvei), nesting material (aspen strips, PM90L, Tapvei) and a clear handling tube (10 cm length, 5 cm diameter). Animal experiments were authorized by the National Animal Experiment Board in Finland (Eläinkoelautakunta, ELLA; license numbers: ESAVI/1172/04.10.07/2018 and ESAVI/1218/2021).

### Tracing

The mice were anesthetized with a mixture of isoflurane (4% for induction, 0.5 – 2% for maintenance; Vetflurane, Virbac, Carros, France) and air (flow rate 0.8 – 1 l/min), after which they were placed into a stereotaxic frame (Kopf Instruments, Tujunga, CA, USA). Before opening the incision on the scalp, iodopovidone was applied to the surgical region, and lubricative eye gel was applied to prevent corneal damage.

For the anterograde tracing studies, stereotaxic coordinates AP: −3.3 ML: ± 0.3 DV: −4.5 mm relative to bregma were used for the VTA, based on the Mouse Brain Atlas (Franklin and Paxinos, 2007) and verified with dye injections. A unilateral injection of 200 nl of AAV2/8-Cag-Flex-Myr-eGFP viral construct (lot 4×1012 genome copies/ml; AAV 319 lot, Neurophotonics Center, CERVO Brain Research Centre, Quebec, Canada) was made with a flow rate of 0.1 µl/min. A precision pump system (KD Scientific, Holliston, MA, USA) was used to control the injection rate.

For the retrograde tracing, the following stereotaxic coordinates were used (mm relative to bregma): for the BNST, AP: −0.1 ML: ±0.9 DV: −4.2; for the CeA, AP: −1.5 ML: ±2.6 DV: −4.7; for the LH, AP: −1.4 ML: ±1.1 DV: −5.3; for the PVT, AP: −1.5 ML: ±0.1 DV: −3.2; for the VP, AP: +0.6 ML: ±1.3 DV: −5.4. For each animal, a 600- nl unilateral injection of 1:1 mixture of green retrobeads (1:10 dilution in dH_2_O, Lumafluor Inc., Durham, NC, USA) and pAAV2-hSyn-DIO-EGFP retrograde virus construct (7.6×10^12^ genome copies/ml; 50457-AAVrg, Addgene, Watertown, MA, USA) was made with a flow rate of 1.0 µl/min. Retrobeads were used only for marking the injection site and were not considered in the backtracing analysis. After the surgeries, the mice were administered 5 mg/kg carprofen (s.c. Norocarp Vet 50 mg/ml, Norbrook Laboratories Ltd, Newry, Northern Ireland) for postoperative analgesia, and were let to recover from anesthesia in a 37°C incubator until ambulatory. A recovery period of at least 3 weeks was allowed before further procedures.

Mice were anaesthetized with pentobarbital (200 mg/kg i.p., Mebunat, Orion Pharma, Espoo, Finland) and perfused transcardially with cold 1xPBS solution followed by 4% paraformaldehyde solution. After overnighting in the same fixative solution, brains were transferred to 30% sucrose until sinking (at least 48 h). The brains were then frozen on dry ice and stored at −80°C until sectioned. For the anterograde tracing, 80-μm coronal sections were cut throughout the brain, and for the retrograde tracing, 40-μm coronal sections from the injection site and the VTA were cut with a cryostat (CM3050S, Leica Biosystems, Wetzlar, Germany).

To enhance the fluorescence signal of the antero-traced axons, anti-GFP immunostaining was performed. The sections were washed at room temperature in 1x PBS (5 min, 3 times), and blocked with 1% bovine serum albumin (BSA; Sigma Aldrich, Saint Louis, MO, USA) with 0.3% Triton X-100 (BDH Laboratory Supplies, Poole, UK) in 1x PBS for 1 h at room temperature. The sections were then incubated with the primary antibody (chicken anti-GFP 1:800 in blocking solution; ab13970, Abcam, Cambridge, UK) overnight at 4°C, washed with 1x PBS (5 min, 3 times) and incubated with the secondary antibody (goat anti-chicken with Alexa Fluor 488 1:800 in blocking solution; ab150169, Abcam) for 2 h at room temperature. Sections were then washed once more with 1x PBS (5 min, 3 times), mounted on microscope slides, and coverslips were applied with Vectashield mounting medium (Vector Laboratories, Burlingame, CA, USA).

Imaging was performed with Zeiss Axio Imager Z2 (Zeiss AG, Oberkochen, Germany) with 10x air objective in tiles mode. The injection site was considered as successful if 90 % of GFP-positive infected cell bodies were located within the VTA area. The Sst nature of the infected cells was also confirmed by the red inbuilt signal of tdTomato. Anterograde targets of the projecting cells were considered as positive by having maximum brightness due to GFP-expressing axonal arborization. The conclusion about the constancy of the projection target was made according to hierarchical clustering results produced by *hclust* function implemented in R programming environment (https://www.rdocumentation.org/packages/stats/versions/3.6.2/topics/hclust). The script and the source data to reproduce the clustering can be found here: https://github.com/eLinanin/Anterograde_heatmap.git.

Retrogradely traced neurons within the VTA were counted as positive by having clear co-expression of the inbuilt tdTomato signal and the GFP signal caused by the viral infection.

### Electrophysiology and single-cell qPCR

To define electrophysiological and molecular profiles of the VTA projecting Sst neurons we combined backtracing with current-clamp recordings and single-cell qPCR. Backtracing was performed as described above (see Tracing section) on P60-P90 old Sst-IRES-Cre animals of both sexes. Electrophysiology was done after at least 1 month of recovery as described previously (Nagaeva et al., 2020) with some modifications. Shortly, mice were perfused with ice-cold constantly oxygenated NMDG-based cutting solution. Coronal VTA sections of 225 µm thickness were cut in the same solution using vibratome HM650V (Thermo Scientific, Waltham, MA, USA) and transferred to the constantly oxygenated 33°C HEPES-ACSF solution for 15 min recovery and then remained in the same solution at room temperature until the end of the experiment (∼ 3 h). For the injection site verification, sections were cut in the same way as the VTA sections and checked immediately with the epifluorescence microscope BX51WI (Olympus, Tokyo, Japan).

Electrophysiological registration of the firing patterns was performed in the same conditions as previously described (Nagaeva et al., 2020) to allow data alignment. Backtraced Sst GFP-positive cells were identified with an epifluorescence microscope BX51WI and sCMOS camera Andor Zyla 5.5 (Oxford Instruments, Oxford, UK). Whole-cell current-clamp recordings were made using 3–5 MΩ borosilicate glass electrodes filled with 1-2 µL of intracellular solution (IS) (containing in mM): 130 K-gluconate, 6 NaCl, 10 HEPES, 0.5 EGTA, 4 Na2-ATP, 0.35 Na-GTP, 8 Na2-phosphocreatine (pH adjusted to 7.2 with KOH, osmolarity ∼285 mOsm). 1 U/µL TaKaRa RNAse inhibitor (Takara, Shiga, Japan) was added before each experiment. Liquid junction potential (+12 mV) was not corrected during recordings.

All electrophysiological experiments were made with a Multiclamp 700B amplifier (Molecular Devices, USA) filtered at 2 kHz, and recorded with a 10 kHz sampling rate using pClamp 10 software (Molecular Devices). After achieving whole-cell configuration in voltage-clamp mode (−70 mV), cell capacitance was determined by the ‘Membrane Test’ feature of the Clampfit software and the amplifier was then switched to current-clamp mode. Depolarized cells with RMP higher than −50 mV were excluded. For measuring passive and active membrane properties, neurons were injected with 800-ms current steps with 10 pA increments, resulting in the membrane voltage fluctuation from −120 mV to the saturated level of firing. In addition, a shorter protocol from 0 pA with 1 pA increment was applied for better identification of the action potential threshold and shape. All recordings were performed with intact GABAergic and glutamatergic transmission (i.e. no pharmacological agents were added to the aCSF).

#### Extraction of mRNA

After firing pattern registration, the cell body was immediately sucked into the glass electrode with negative pressure and expelled onto a 1.1 µl drop of the ice-cold lysis buffer (0.1% Triton X-100, 2 U/µl TaKaRa RNAse inhibitor, 0.5 µM Oligo-dT_30_ primer, 11.5 mM dithiothreitol and 2.3 mM of dNTP mix) placed on the wall of cold RNAse-free PCR tube, spun down and immediately frozen in dry ice. All samples were stored at −80°C until the reverse transcription step.

#### Reverse transcription (RT) and pre-amplification

Frozen samples were thawed at 72°C for 3 min. 2 µl of RT mix (1x SuperScript IV buffer, 18 U/µl SuperScript IV reverse transcriptase (Thermofisher Scientific, USA), 3.6 µM TSO-chimera primer, 6 mM MgCl2, 0.8 M betaine and 1.5 U/µl TaKaRa RNAse inhibitor) were added to each sample, mixed gently, spun down and then incubated in the thermocycler (52°C − 10 min, [60°C − 1 min, 52°C − 1 min] x 10 times, 80°C − 20 min). For further cDNA amplification, 20 µl of PCR mix [1xPlatinum SuperFi II master mix, 0.5 µM PCR1 primer, 111 nM dNTP mix] were gently pipetted to each sample, which then underwent the following thermocycler program (98°C − 30 s, [98°C − 10 s, 60°C − 11 s, 72°C − 6 min] x 25 times, 72°C − 5 min).

#### qPCR analysis

The synthesized cDNA samples were diluted to the concentration range 100-300 ng/μl with sterile DNAse free water. Samples were then amplified with PowerUp™ SYBR™ Green Master Mix (Applied Biosystems, USA) following the instructions of the manufacturer. Quantitative PCR (qPCR) was performed using LightCycler® 480 II system (Roche Diagnostics, Switzerland) with the following conditions: UDG (uracil-DNA glycosylase) activation 2 min at 50°C, pre-incubation 2 min at 95°C, followed by amplification steps 40 cycles of 15 s at 95°C and 1 min at 60°C with the melting curve. Expression of the following genes was determined: *Sst*, *Th*, *Egfp*, *Nos1, Ddc, Gad1*, *Gad2*, *Slc17a6* (*Vglut2*), *Slc32a1* (*Vgat*), and *Slc6a3* (*Dat*). The primer sequences were obtained from the PrimerBank (https://pga.mgh.harvard.edu/primerbank/index.html). Primers were ordered from Metabion (Metabion International AG, Germany) and actual sequences are listed in the table below this section. Every run included a negative control, a positive control (bulk cDNA from mouse visual cortex) and a blank (H_2_O). The qPCR reactions were performed in triplicate for each sample, and their averages were used for Ct values. The Ct values of EGFP were used to normalize expression by the delta Ct method as the EGFP was present in both backtraced and DA neurons from Th-EGFP mice. The relative expression was calculated based on the delta Ct values. Resulting values were then normalized with log10 transformation for the clustering procedure performed as described in the “Tracing” part. The script for the clustering can be found here: https://github.com/eLinanin/Fig4c_qPCR_heatmap.git.

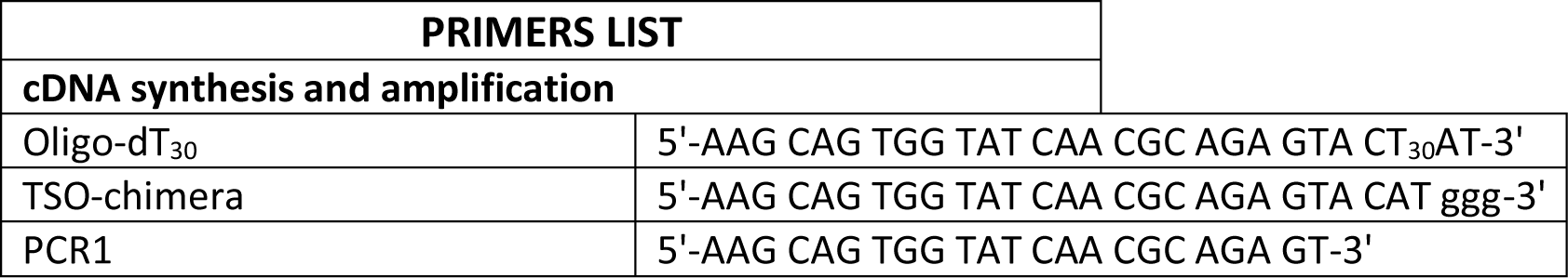

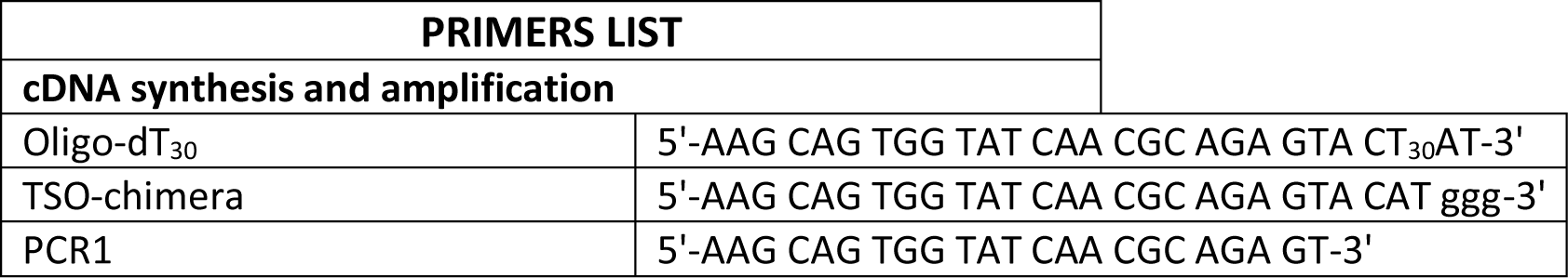

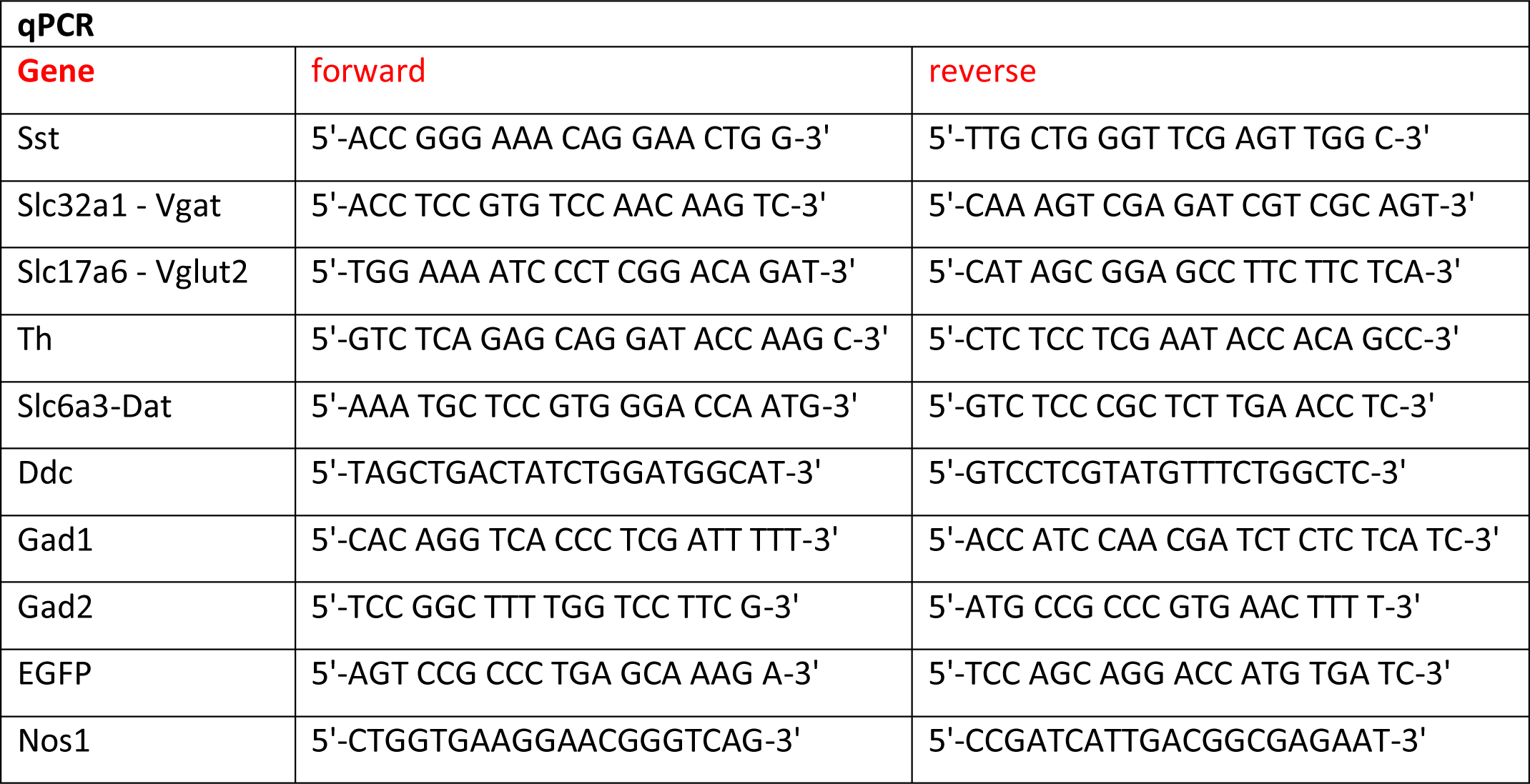

#### Caspase experiments

For the caspase manipulations, Sst-tdTomato male and female mice of the age P60-P75 (on the injection day) were used. All the injection procedures were similar to those described in the “Tracing” part of the Methods. Slightly modified coordinates for the VTA were used, aiming at the anterolateral Sst sub-population (Nagaeva et al., 2020) and based on the results of the retrograde tracing to hit the majority of the projecting VTA Sst neurons (mm relative to bregma): AP: −3.1, ML: ±0.7, DV: −4.6. 600-nl bilateral injections of 1:1 mixture of AAV1/2-DIO-taCasp3-TEV (3×10^12^ genome copies/ml, a kind gift from professor William Wisden) and AAV2/8-CAG-Flex-Myr-eGFP (4×10^12^ genome copies/ml; Neurophotonics Center, CERVO Brain Research Centre, Quebec, Canada) viral constructs, or 1:1 dH_2_0 dilution of AAV2/8-CAG-Flex-Myr-eGFP for the controls, were made with a flow rate of 0.1 µl/min. After surgery, mice had a recovery period of 4 weeks before the behavioural tests.

Confirmation of the deletion of VTA Sst neurons was made after the end of all behavioural experiments. Mice were perfused as described in the “Tracing” methods section. Forty-µm-thick coronal sections containing the VTA area were made and imaged with the epifluorescence microscope. Numbers of the eGFP- expressing cells were counted in the control and caspase groups and compared. The caspase brains where the injection site did not reach the VTA, were too lateral, or too medial were excluded from the analysis (4 mice from cohort 1; 2 from cohort 2, and 5 mice from the “Intellicage cohort”). The subjective exclusion was performed by two independent researchers.

### Behavioural experiments

#### General description

17 (9 male, 8 female) VTA^Sst-^ and 21 (10 male, 11 female) VTA^Sst+^ control mice underwent a battery of behavioural tests to study reward- and anxiety-related behaviours. Experiments were performed in the morning between 7 am and 1 pm. Mice were always habituated to the testing room 45 – 60 min prior to the experiment. Arenas were cleaned thoroughly with water between animals. Researchers conducting thee xperiments were blinded to the treatment group of the animals. Mice were initially group-housed, but then single-housed (including 3 days of habituation) for conducting the novelty-suppressed feeding, sucrose preference, running wheel activity and morphine sensitization tests (see the schematic timeline below, W – weeks). To reduce stress from the injection, mice underwent a 5-day habituation routine prior to morphine injections as described before (Elsilä et al., 2022).

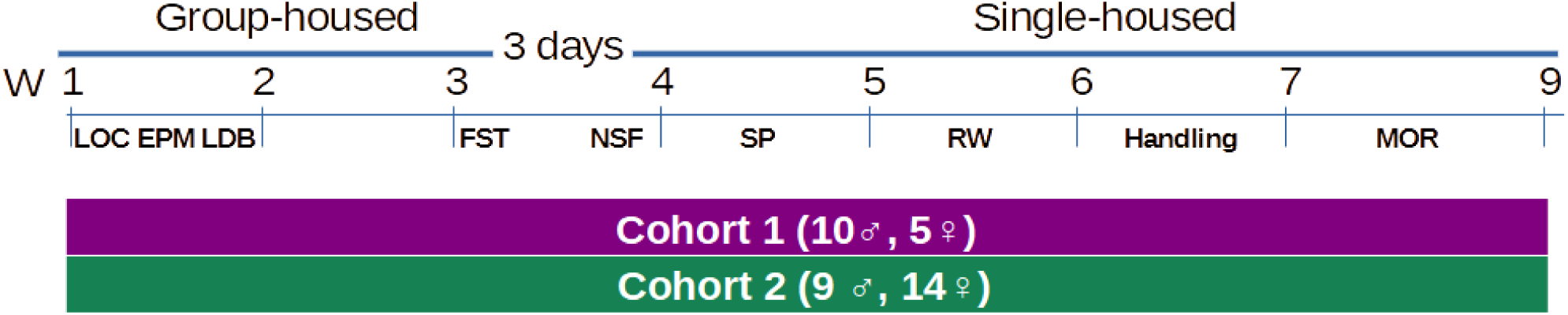

#### Novelty-induced locomotor activity (LOC) and morphine sensitization (MOR)

The experiment was performed as described previously (Vashchinkina et al., 2012). Shortly, mice were released one by one in the novel open arena (36 x 19 x 20 cm). Distance moved was recorded during 60 min with EthoVision XT 10 tracking equipment (Noldus, Netherlands). Illumination in the room was approximately 50 lux.

For **morphine sensitization**, each mouse was habituated to the arena for 60 min after which it received a morphine injection (20 mg/kg, i.p.) and was immediately placed back in the arena. Mice remained in the arena for 3 h, and locomotor behaviour was monitored using EthoVision (induction day). After the testing, the mice were put back in their home cage. Mice were challenged with a morphine injection 7 days later, and the experiment was repeated in the same context (challenge day). Sensitization to the morphine-enhanced locomotor activity was calculated as the difference of distance moved on the challenge day minus distance moved on the induction day. Two male caspase-treated mice were excluded from the analysis due to the problems with the first (induction) morphine injection. Exclusion did not change the statistical outcome.

#### Elevated plus maze (EPM)

The elevated plus maze was made out of grey plastic and consisted of a central platform (5 x 5 cm), from which two open arms and two enclosed arms (5 x 40 x 20 cm) extended at an elevation of 50 cm from the floor (Lister R.G., 1987). The light intensity of the closed arms was 10 lux and open arm 200 lux. The mouse was placed on the central platform facing a closed arm and allowed free exploration of the maze for 5 min. Distance moved and time spent in different arms was recorded with EthoVision. Time spent in the open arm was calculated as the percentage of the total time spent in all arms.

#### Light-dark box (LDB)

The test was performed in an open-field arena (43.2 x 43.2 x 30.5 cm, ENV-515, Med Associates Inc., St. Albans, VT) equipped with infrared light transmitters and sensors detecting horizontal and vertical activity. The dark insert (non-transparent for visible light) was used to divide the arena into two equally sized compartments. An open door (width of 5.5cm and height of 7 cm) in the wall of the insert allowed the animal to freely move between compartments. Illumination in the light compartment from bright ceiling lights was ∼ 200 lm. The animal was released in the door, head facing the dark compartment and allowed to explore the arena for 5 min. Distance moved and time spent in different compartments were recorded by the system. Time spent in the light compartment was calculated as the percentage of the total time spent in both compartments.

#### Forced swim test (FST)

The mouse was placed for 6 min in a glass beaker (diameter 15 cm, height 25 cm) filled with 3 L of water at 23 ± 1°C (Procaccini et al., 2011). Three visually isolated mice were recorded simultaneously using a digital video camera. Latency to the first immobility and time of total immobility (i.e. passive floating, when an animal was motionless and only doing a slight movement with a tail or one hind limb, in contrast to struggling, climbing or swimming with all four paws) were measured manually in 2-min intervals by a blinded researcher.

#### Novelty-suppressed feeding (NSF)

Mice were single-housed and food-restricted for 14 h before testing to motivate food-seeking behaviour. Water was given *ad libitum*. Mice were tested in an open arena (50 x 50 x 28 cm) at an illumination of 200 lux. A lid from a 50-mL falcon tube was placed in the middle of the arena that contained a small amount of moist food. The mouse was released next to the wall, and its behaviour was recorded using EthoVision. Latency until the mouse started eating (i.e. eating for more than 5 s) was monitored manually by a researcher, after which the trial was terminated (maximum cut-off point 10 min). The mouse was returned to its home cage, where it was again presented with a small amount of moist food. Latency until first in-cage eating was monitored as previously. This was done to account for possible differences in hunger between animals.

#### Sucrose preference test (SP)

Two-bottle choice sucrose preference test was carried out for 7 days (Lainiola et al., 2019). Mice were single-housed in larger IVC cages allowing the use of two drinking bottles. Drinking bottles were weighted daily at 10:00 am to monitor consumption. The position of the water and sucrose bottles was changed every day to avoid the development of side preference. Mouse body weights were measured before and after the start of the experiment. Sucrose concentrations were based on an earlier study analyzing sucrose preference in C57 mice (Sclafani, 2006) and were increased as follows: 0.1% (two days), 0.5% (two days) and 1% sucrose (three days). The average was taken for each sucrose concentration. Sucrose preference was calculated as a percentage of sucrose consumption out of the total fluid intake.

#### Circadian rhythm of wheel running (RW) activity

To measure voluntary wheel running activity, free-running wheels (MedAssociates Inc.,) were placed in the IVC cage of a single-housed mouse. The rotation of the wheel by the mouse was transmitted as a digital signal wirelessly to a hub and recorded on the Wheel Manager software. Data was exported every morning, and running wheels were checked for proper functioning. Voluntary running wheel activity was followed for 3 days, of which the first day was considered as habituation followed by two days of basal activity. Ten mice, which did not use the running wheel and ran <100 rotations over a course of 3 days, were excluded from the analysis.

#### IntelliCage (IC)

A separate “Intellicage cohort” of 15 males and 16 females was subcutaneously injected with RFID transponders (Planet ID GmbH, Germany) for individual identification. The IntelliCage by NewBehaviour (TSE Systems, Germany) is an apparatus designed to fit inside a large cage (610 x 435 x 215 mm, Tecniplast 2000P). The apparatus itself provides four recording chambers that fit into the corners of the housing cage. Access into the chambers was provided via a tubular antenna (50 mm outer and 30 mm inner diameter) reading the transponder codes. The chamber contains two openings of 13 mm in diameter (one on the left, one on the right), which gave access to drinking bottles. These openings are crossed by photo beams recording nose-pokes of the mice and the holes can be closed by motorized doors. Four triangular red shelters (Tecniplast, Buguggiate, Italy) were placed in the middle of the IntelliCage and used as sleeping quarters and as a stand to reach the food. The floor was covered with a thick (2-3 cm) layer of bedding. The IntelliCage was controlled by a computer with dedicated software (IntelliCagePlus), executing preprogrammed experimental schedules and registering the number and duration of visits to the corner chambers, nose-pokes to the door openings and lickings as behavioural measures for each mouse. To randomize treatment groups and allow non-competitive access to the corners mice were housed in 4 Intellicages in balanced groups of the same sex (e.g. 4 control + 4 caspase). The mice were group-housed in these groups from weaning (cage type Tecniplast Green Line GR900), at least 10 weeks before the start of Intellicage experiments. All tests in the Intellicage system were done in the order they are listed below on the consecutive days without taking mice out from the cages except as on two cleaning days.

#### Free adaptation

At the beginning of the test, the mice were released in the IntelliCage during the light phase at 9 am with all doors open allowing unlimited access to the water bottles (free adaptation). Animals were allowed to explore the new environment for 3 consecutive days. The exploratory, locomotor and circadian activities were measured as a number of corner visits or as nose-pokes to the water bottles per hour for each day separately (day 1 – adaptation phase, day 2 and 3 – basic activity). Similarly, drinking behaviour was measured as the number of licks/h.

#### Adaptation to nose-poke

On the fourth day, all doors were closed at the beginning of the experiment and mice were required to poke into closed gates to reach drinking tubes. Only the first nose-poke of the visit opened the door for 5 seconds (pre-defined time). Animals had to start a new visit in order to get access to water again. This rule was the same in all experiments requiring nose-poking.

#### Saccharine preference

In this task, all four corners operated the same way, 24 h per day: doors opened spontaneously for a 7-s drinking period on the entry to a corner. Each corner contained a bottle of saccharine on one side and a bottle of water on the other. High and low saccharine concentrations were chosen based on previous research (Pijlman et al., 2003; Sclafani et al., 2010). Every day, saccharine and water sides were alternated to exclude side preference. During the first three days, mice were suggested to choose between two low saccharine concentrations 0.01% (**S1**) and 0.03% (**S2**) assigned to two opposite corners each. During the next three days, the lowest 0.01% (**S1**) concentration was exchanged for the highest 0.3% (**S3**) one and the order of the corners was changed as well. For a better understanding of the schedule, below there is a simplified scheme with the corners and sides assignments.

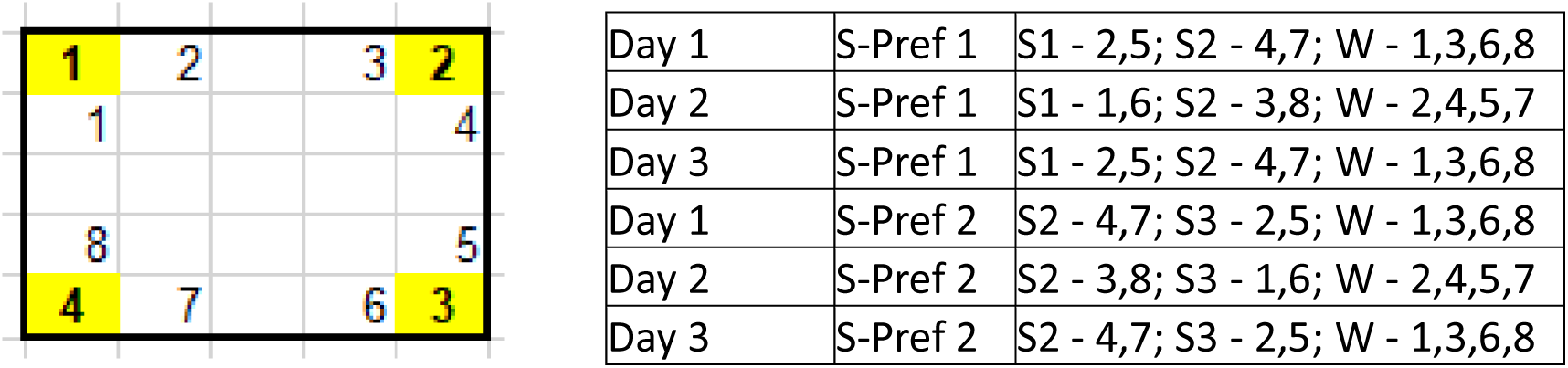

The preference score was calculated as a percentage of the number of licks to a certain liquid (two saccharine liquids of different concentrations and two water liquids in corresponding saccharine corners) from the total number of licks during the last two days of each session.

#### Delay discounting

In this experiment, all four corners were accessible to all animals and contained 0.3% saccharine liquid (**S3**) on one side of the corner and water on the other side. The order of the bottles was as follows (see the picture above): S3 - 1,3,6,8; W - 2,4,5,7. On Day 0 both doors to the water and saccharine opened simultaneously for 7 s upon entry to the corner. On Day 1, saccharine door opened with a 0.5-s delay, while the water bottle door opened immediately. The next day, the delay before the opening of the saccharine door increased to 1 s and then for 1 s every next 24 h. After 4 days, this resulted in a delay of 5 s. A saccharinepreference score was calculated as a percentage of the lick number to saccharine bottles from a total lick number (saccharine + water).

#### Saccharine extinction and avoidance

In these two tasks, the set-up was very similar in general and differed only in the third phase. Water was always available on the entry in two corners for all animals. Saccharine 0.3% (**S3**) was available with a rule in one corner for 4 specific mice (2 caspase+2 controls) and in another corner for the remaining 4 mice to avoid competition. The rule was as follows: to get the saccharine mice had to nose-poke in one of the side doors, it triggered the LED light above this door for 1.5 s and then it opened after an extra 0.5-s delay for 5 s. Mice had to repeat the sequence in order to get additional access to the saccharine. In Phase 1 (33 h), mice were learning the rule; in Phase 2 (48 h) mice were adapted to the rule and the basic activity was measured; in Phase 3 (38 h) saccharine bottles were emptied for the “Extinction” experiment. In the “Avoidance” experiment mice went through the same sequence of events, but in Phase 1 saccharine bottles were back, and the order of the corners was changed, so they had to learn new rules (re-learning). In Phase 3 (Avoidance), mice received 0.2-bar air puffs in the saccharine door with a 25% probability. The activity was measured during the whole experiment as a number of nose-pokes/h. Similarly, drinking behaviour was measured as licks/h.

#### Fear conditioning (FC)

For these experiments, the same “Intellicage cohort” of mice was used. Animals were single-housed after the Intellicage experiments and given two weeks of adaptation prior to the FC. The FC protocol was based on previously published studies (Mennesson et al., 2020) with few modifications and consisted of three phases: acquisition, context test and cue test. Shortly, mice were placed one by one in the test chamber (Video fear conditioning, Med Associates Inc.) for fear *acquisition and conditioning*. After 120-s of free exploration, a 30-s 5 kHz 90-dB cue tone sounded from the wall-mounted speakers, co-terminated with a 2-s scrambled 0.6-mA shock through the grid floor. Cue-shock pairs were repeated twice again with 90 and 60 s inter-trial intervals, after that the session was finished with 120 s of free exploration. Chamber light, near-infrared light and a fan were on during all phases. After each mouse the chamber was cleaned with water.

Nine days later, the mice were tested in the same chambers for the *context*-induced retrieval of the fear memory. For that, the mouse was placed in the same testing chamber with identical conditions as before, except for no cue tones or shocks, for 300 s of free exploration.

Five hours after the context test, *cue*-induced retrieval of the fear memory was assessed in the conditioning chamber with the floor and wall material and the shape of the chamber changed to exclude the context component. After 120 s of free exploration, the mouse was introduced to twenty 30-s cue tones (identical to those used during conditioning) separated by 5-s inter-tone intervals. After each mouse, the chamber was cleaned with 70% ethanol.

Freezing time and the number of freezing episodes were automatically analyzed by Video Freeze Software (Med Associates Inc.) separately for each component of the test phases. Two female mice had missing data for the acquisition phase, but they still received the foot shock and were included in the context- and cue-induced memory retrieval data analysis.

#### Drugs

Morphine hydrochloride (University Pharmacy, Helsinki, Finland) was dissolved in physiological saline (0.9% NaCl) on the day of treatment. Morphine was injected in a volume of 10 mL/kg body weight.

#### Statistics

For behavioural experiments, statistical analysis was done using SPSS (IBM SPSS Statistics, 28.0.0.0) while graphs were drawn with Graphpad Prism 8.1.10. The data were tested for normality and homogeneity of variance using the Kolmogorov- Smirnov and Levene’s tests, respectively. When assumptions were violated, the square root transformation was applied (“elevated plus maze” and the “running wheel”). Statistical analyses of the data were done using univariate and repeated measures two-way ANOVAs unless stated otherwise. In case of significant main effect or interaction, post-hoc tests were performed using multiple comparisons test with Bonferroni correction. The level of significance was set at 0.05. All data are shown as means ± SEM.

## Acknowledgements

The following core facilities were essential for the project: Biomedicum Imaging Unit of the University of Helsinki; Mouse Behavioural Phenotyping Facility of the University of Helsinki, supported by Helsinki Institute of Life Science and Biocenter, Finland; Biostatistics Consulting Service of the University of Helsinki; Canadian Neurophotonics Platform, Québec, Canada. The authors are grateful for the expert technical assistance and fruitful discussion by Heidi Hytönen, Vootele Voikar, Ivan Zubarev, Merja Voutilainen, Mikko Airavaara, Lauren van den Broecke and Laura Martikainen.

## Supplementary Figures

**Figure S1.**
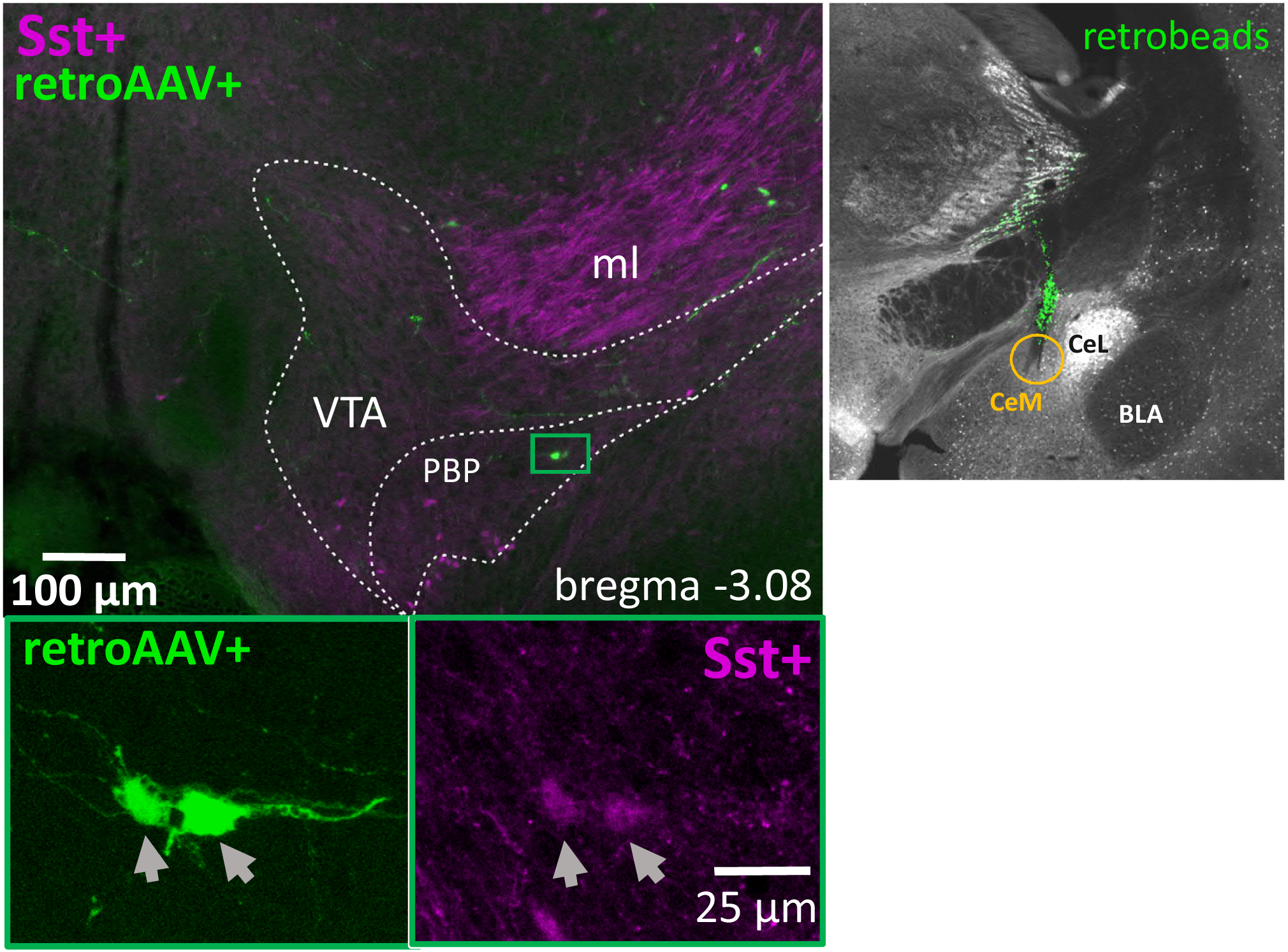
Backtracing from the medial part of the Central Amygdala. Examples of the backtraced neurons in the VTA at the bregma level -3.08 mm in Sst- tdTomato (magenta) mouse. The image on the right shows retrobeads in the injection site (CeM). The yellow circle shows the actual unilateral injection spot. On the top left, the green rectangle shows ipsilaterally traced neurons. The lower panels are the magnified images of the green rectangle. ***BLA*** – basolateral amygdala; ***CeL*** – lateral part of the central amygdala; ***CeM*** – medial part of the central amygdala; ***ml*** – medial lemniscus; ***PBP*** – parabrachial pigmented nucleus of the VTA; ***VTA*** – ventral tegmental area.

**Figure S2.**
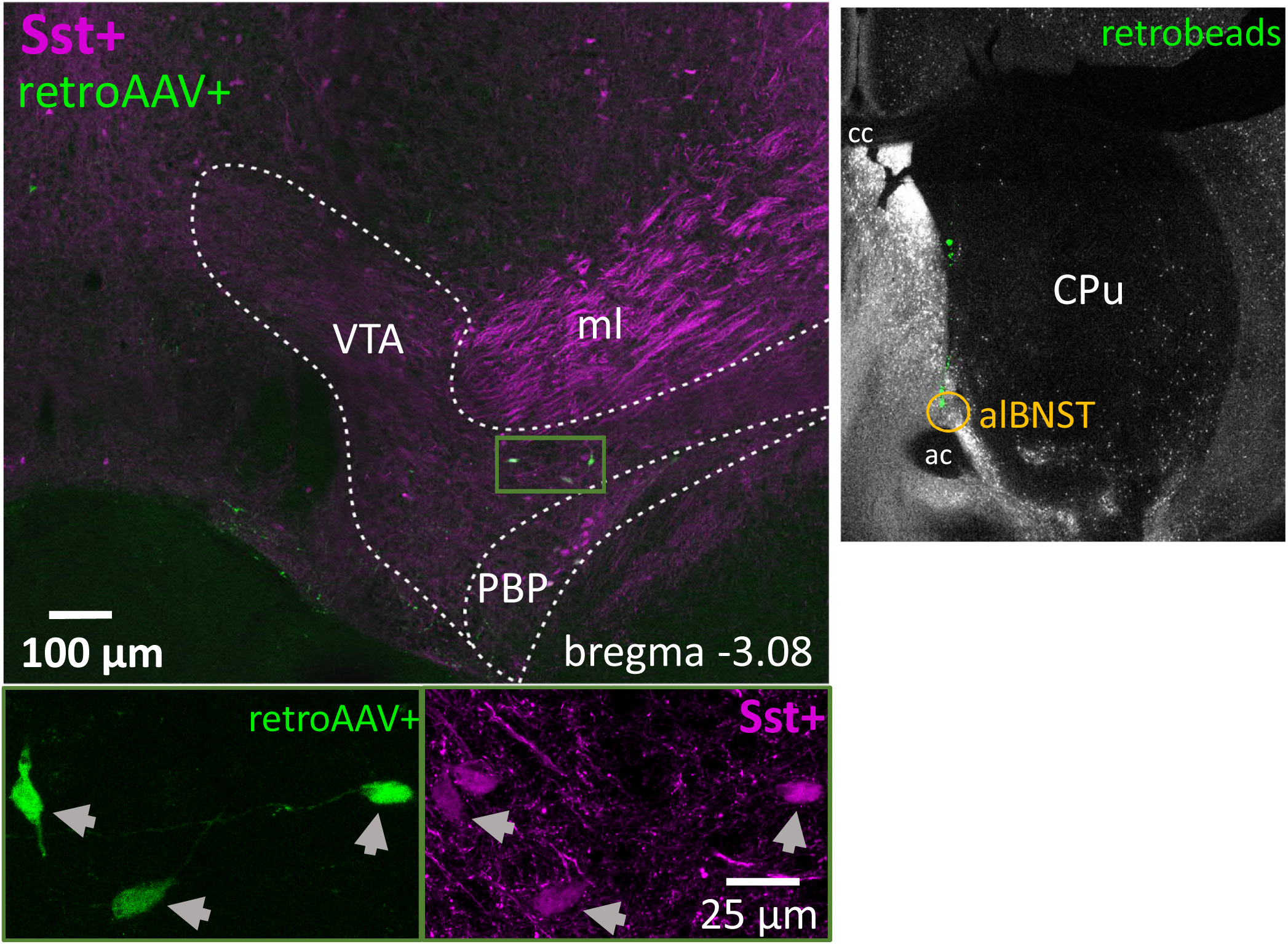
Backtracing from the anterolateral part of the Bed Nucleus of Stria Terminalis. Examples of the backtraced neurons in the VTA at the bregma level -3.08 mm in Sst-tdTomato (magenta) mouse. The image on the right shows retrobeads in the injection site (alBNST). The yellow circle shows the actual injection spot. On the top left, the green rectangle shows ipsilaterally traced neurons. Lower panels are magnified images inside green rectangle split by fluorescent channels. ***ac*** – anterior commissure; ***alBNST*** – bed nucleus of the stria terminalis, antero-lateral part; ***cc*** – corpus callosum; ***CPu***- caudatus-putamen (striatum); ***ml*** – medial lemniscus; ***PBP*** – parabrachial pigmented nucleus of the VTA; ***VTA*** – ventral tegmental area.

**Figure S3.**
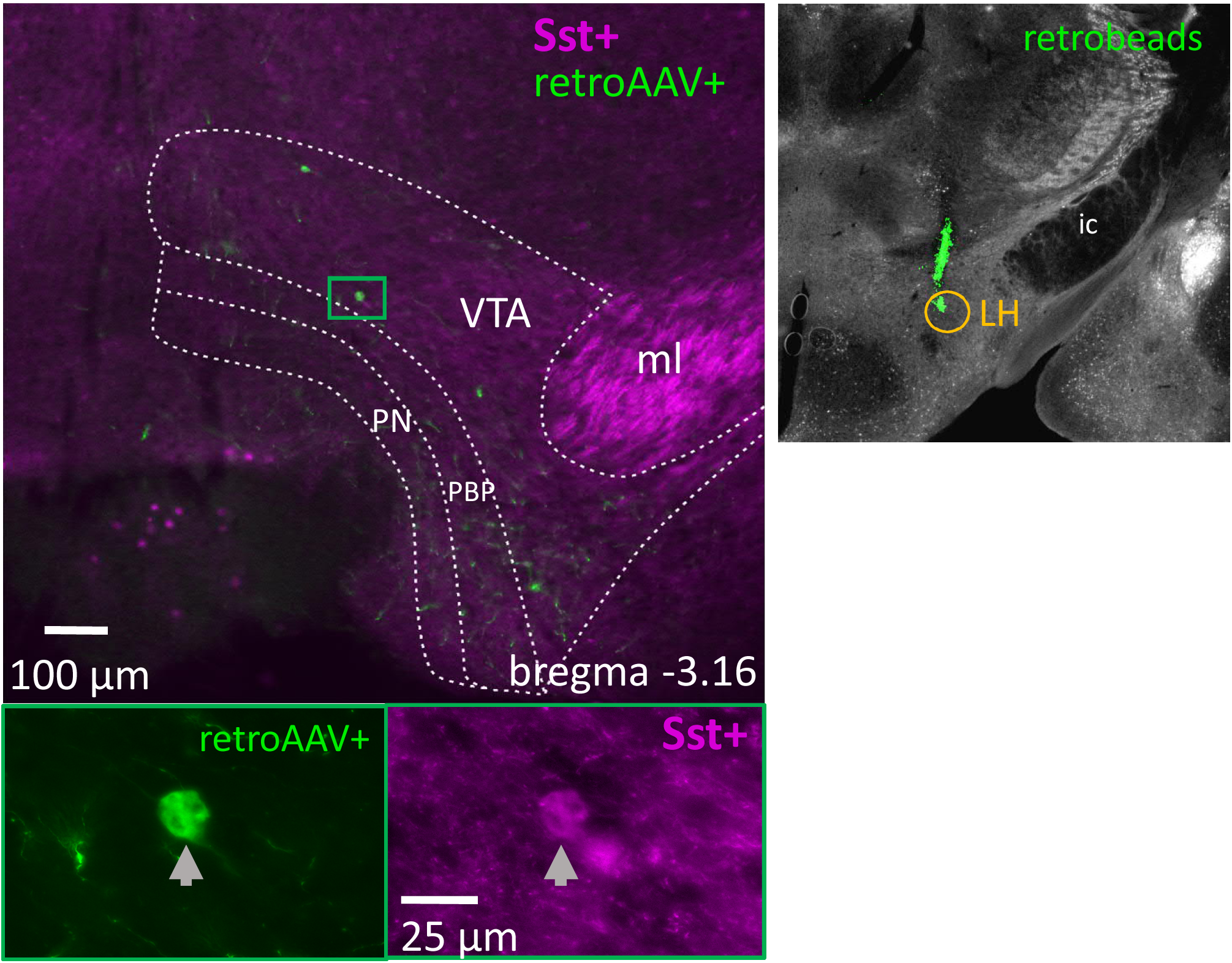
Backtracing from the Lateral Hypothalamus. Examples of the backtraced neurons in the VTA at the bregma level -3.28 mm in Sst-tdTomato (magenta) mouse. The right upper corner shows retrobeads at the injection site (LH). The yellow circle shows the actual unilateral injection spot. On the top left, the green rectangle shows an ipsilaterally traced neuron. Lower panels are magnified images inside the green rectangle split by fluorescent channels. ***ic*** – internal capsule; ***LH*** – lateral hypothalamus; ***ml*** – medial lemniscus; ***PBP*** – parabrachial pigmented nucleus of the VTA; ***PN*** – paranigral nucleus of VTA; ***VTA*** – ventral tegmental area.

**Figure S4.**
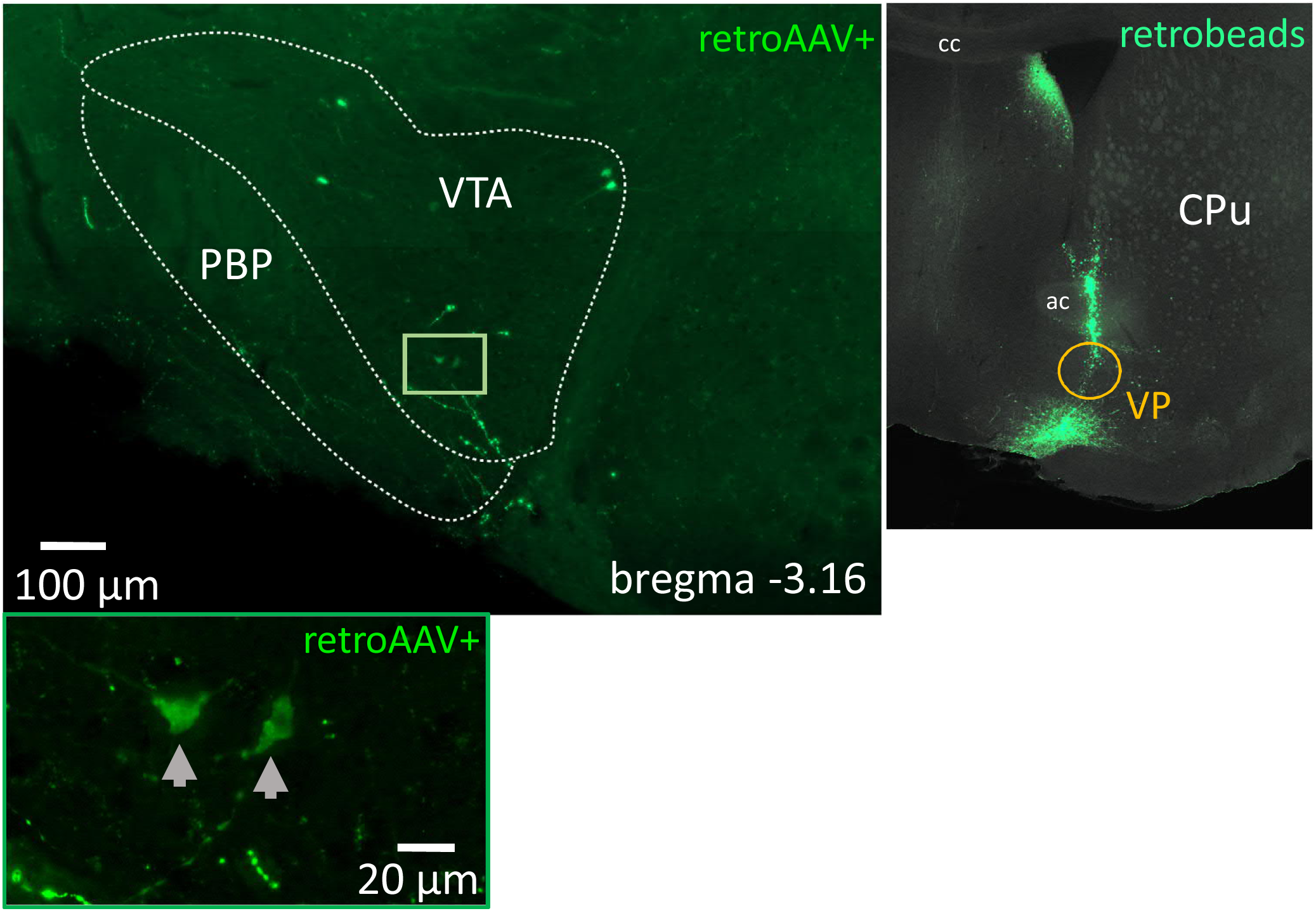
Backtracing from the Ventral Pallidum. Examples of the backtraced neurons in the VTA at the bregma level -3.08 mm in Sst-Cre mouse. The image on the right shows retrobeads in the injection site (VP). The yellow circle shows the actual unilateral injection spot. On the top left, the green rectangle shows ipsilaterally traced neurons. The lower panel is the magnified image of the green rectangle. ***ac*** – anterior commissure; ***cc*** – corpus callosum; ***CPu***- caudatus-putamen (striatum); ***PBP*** – parabrachial pigmented nucleus of the VTA; ***VP*** – ventral pallidum; ***VTA*** – ventral tegmental area.

**Figure S5.**
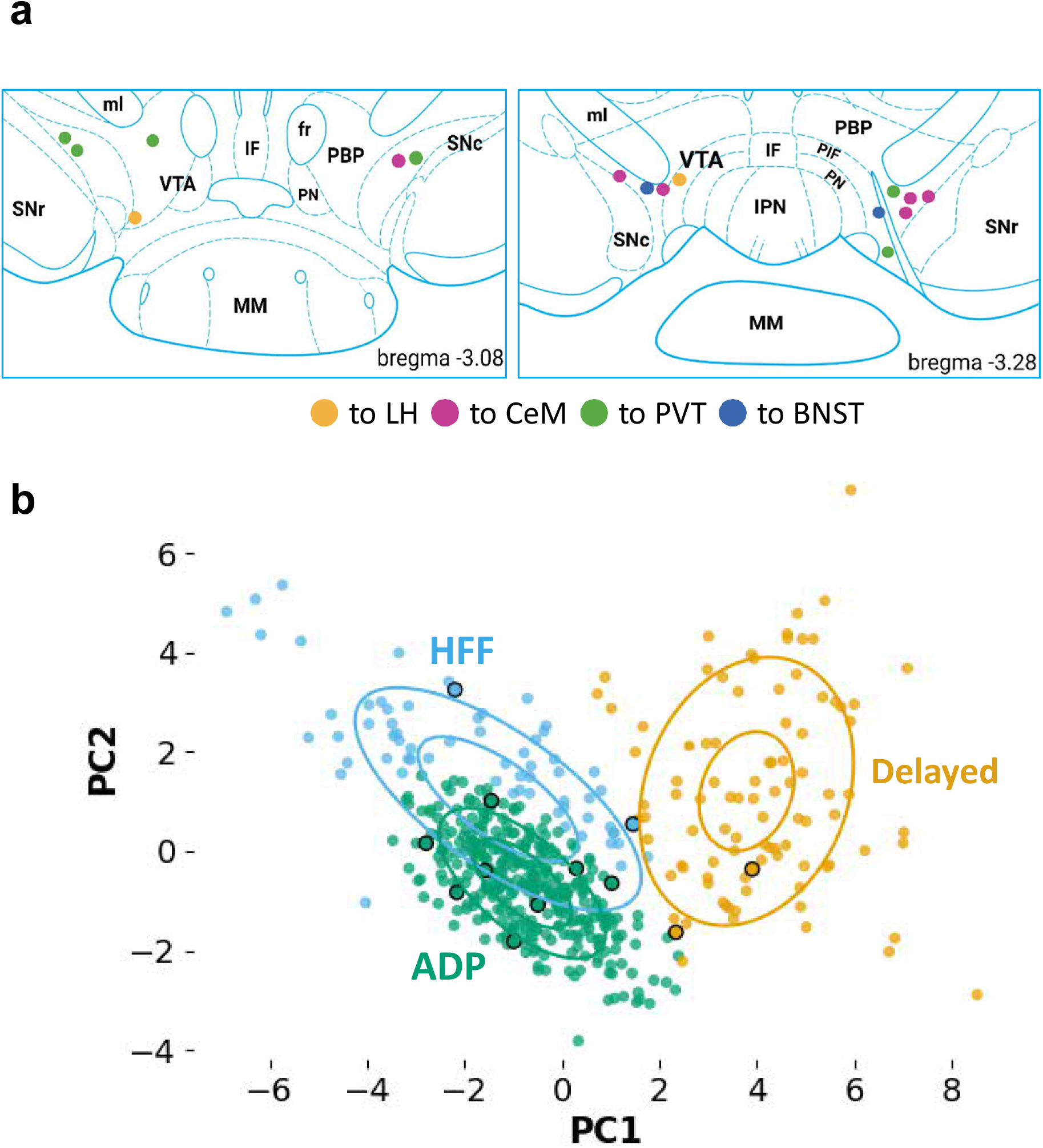
Location of the electrophysiologically recorded VTA Sst neurons projecting to forebrain regions and their electrophysiological subtypes. **a.** None of the backtraced neurons in electrophysiological experiments were found more posterior than the bregma level -3.28 mm, and most of them were located in the lateral nuclei of the VTA. Their projection sites are colour-coded. **b.** Most of the recorded backtraced VTA neurons (black-circled) were assigned to the ADP cluster by unsupervised clustering procedure with a previously published dataset of the VTA Sst neurons as the reference. (The method and clusters were described previously in Nagaeva et el., 2020 – see Fig.3). ***BNST*** – bed nucleus of the stria terminalis; ***CeM*** – central amygdala, medial part; ***LH*** – lateral hypothalamus; ***PVT*** – paraventricular nucleus of the thalamus. ADP – afterdepolarizing, HFF – high-frequency firing.

**Figure S6.**
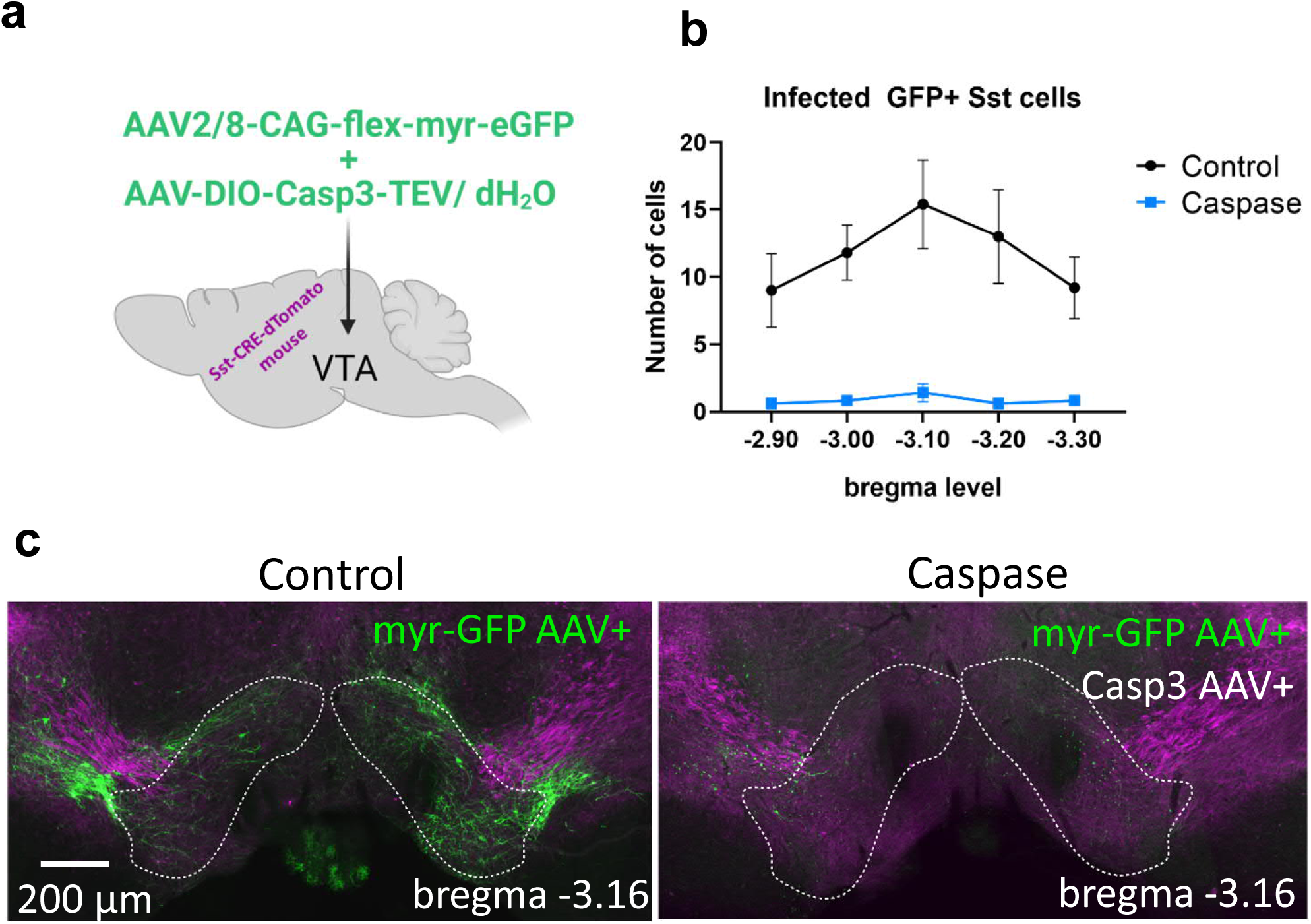
Deletion of the VTA Sst neurons with caspase 3 expressing virus. **a.** Scheme of the bilateral intra-VTA viral injections to Sst-tdTomato (magenta) mice. The control group received only myr-eGFP virus diluted with dH_2_0 to adjust the final volume. **b.** The graph shows an average number of GFP+ Sst cell bodies in the control and caspase-treated animals (n=5 animals per group) depicted per bregma level (X-axis). **c.** Example images of the mouse coronal VTA section from the control (on the left) and caspase group (on the right). The caspase image has almost no infected GFP+ cell bodies in the VTA region (outlined with a white dashed line), showing only sparse GFP+ neurite fragments of the dead Sst neurons.

**Figure S7.**
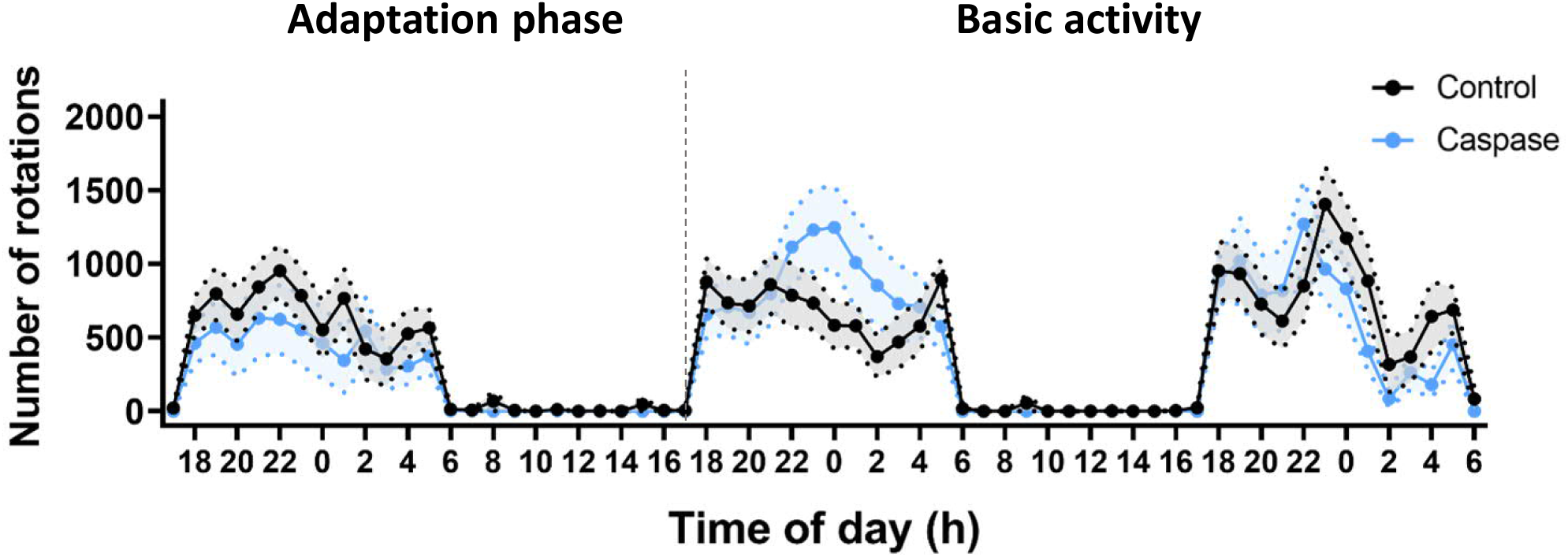
Deletion of VTA Sst neurons did not affect circadian activity in the free-running wheel test (lights on 6-18). Activity of the control (black) and VTA**^Sst^**-caspase (blue) mice. There were no differences in the number of rotations between the treatment groups across three days (treatment: F(1,24)=0.202, p=0.657; treatment x time: F(67,1608)=1.230, p=0.278). Data are shown as means ± SEM.

**Figure S8.**
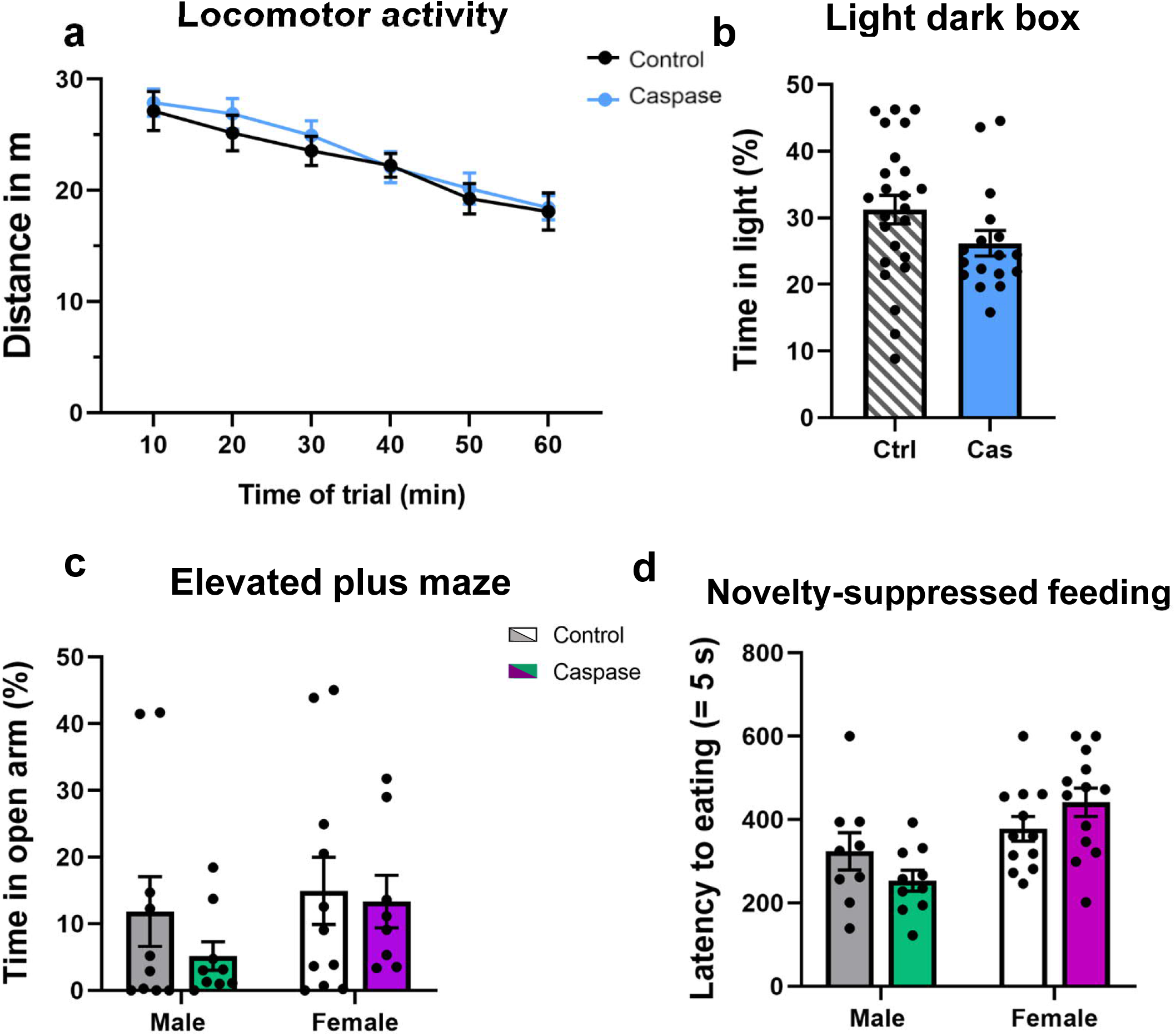
Deletion of Sst neurons in the VTA had no effect on locomotor activity or anxiety-like behaviour. **a.** Locomotor activity in the open arena was not different between the VTA**^Sst^**-caspase and control mice (F(1,34) = 0.265, p = 0.61). **b.** The light-dark box test did not show any difference in percentage of time spent in the light compartment between the treatment groups (F(1,34) = 1.750, p=0.195, sex F(1,34) = 3.957, p= 0.055). **c.** Similarly, percentage of time spent in the open arm measured in the elevated plus maze test was not different. **d.** Latency to start eating in a novel environment did not show a difference, albeit a marginal significance for sex-dependent effects was detected in the VTA^Sst^-caspase mice (treatment x sex: F(1,34) = 3.862, p = 0.058). Data are shown as mean ± SEM.

**Figure S9.**
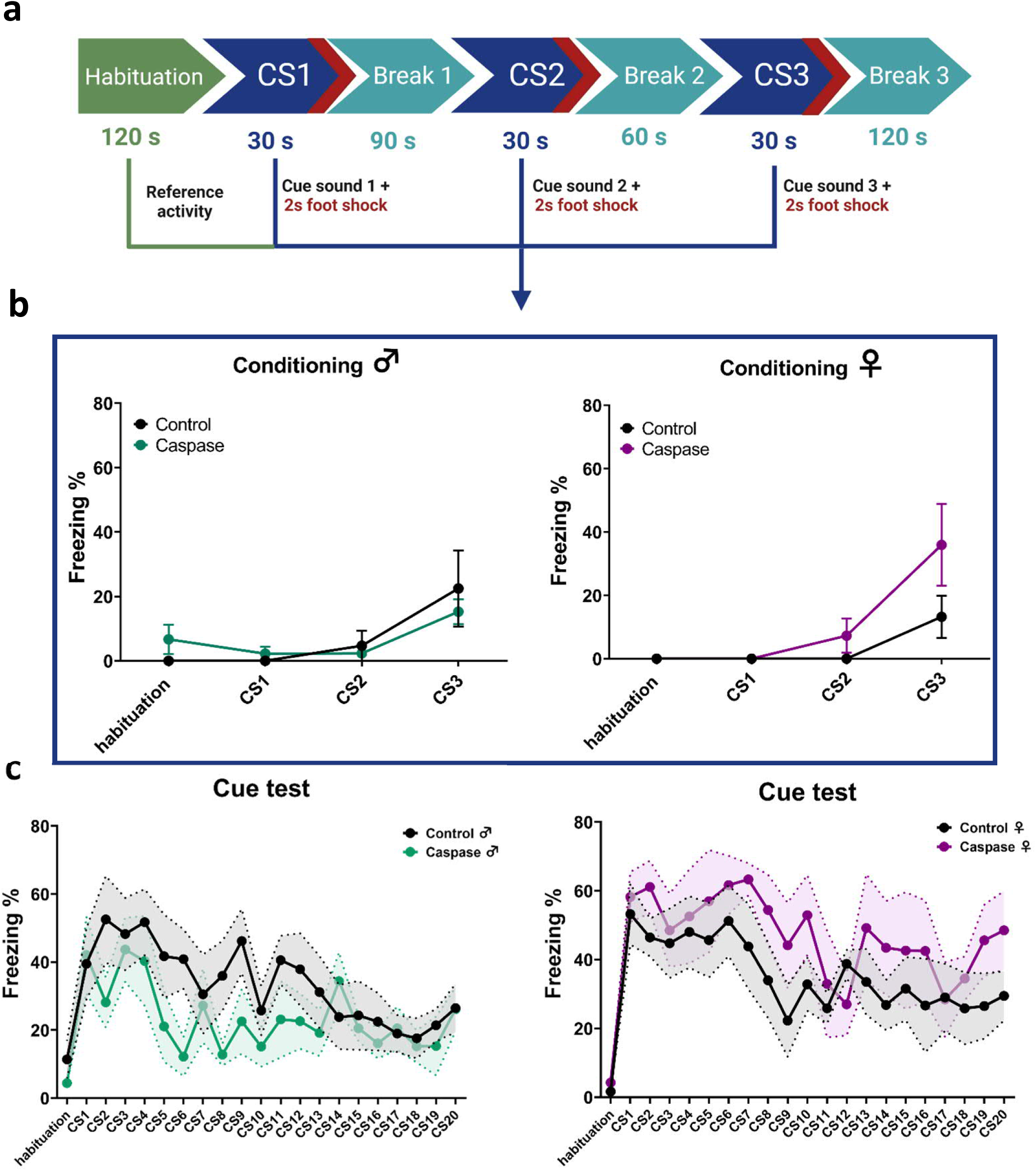
Deletion of the VTA Sst neurons did not affect cue-induced fear processing in Pavlovian fear conditioning. **a.** Protocol of fear conditioning during the acquisition phase. **b.** Graphs show per cent freezing (freeze time/total time) during 30-s cue-sound presentations, co-terminated with 2-s foot shocks. There was no difference in freezing between sexes (F(1,19)=0.019, p=0.891) or between treatments (F(1,19)=2.182, p=0.156) . **c.** Similarly, there were no significant difference in rates of cue-associated fear memory retrieval or extinction (cue x sex x treatment: F(1,420)=1.04, p=0.413).

**Figure S10.**
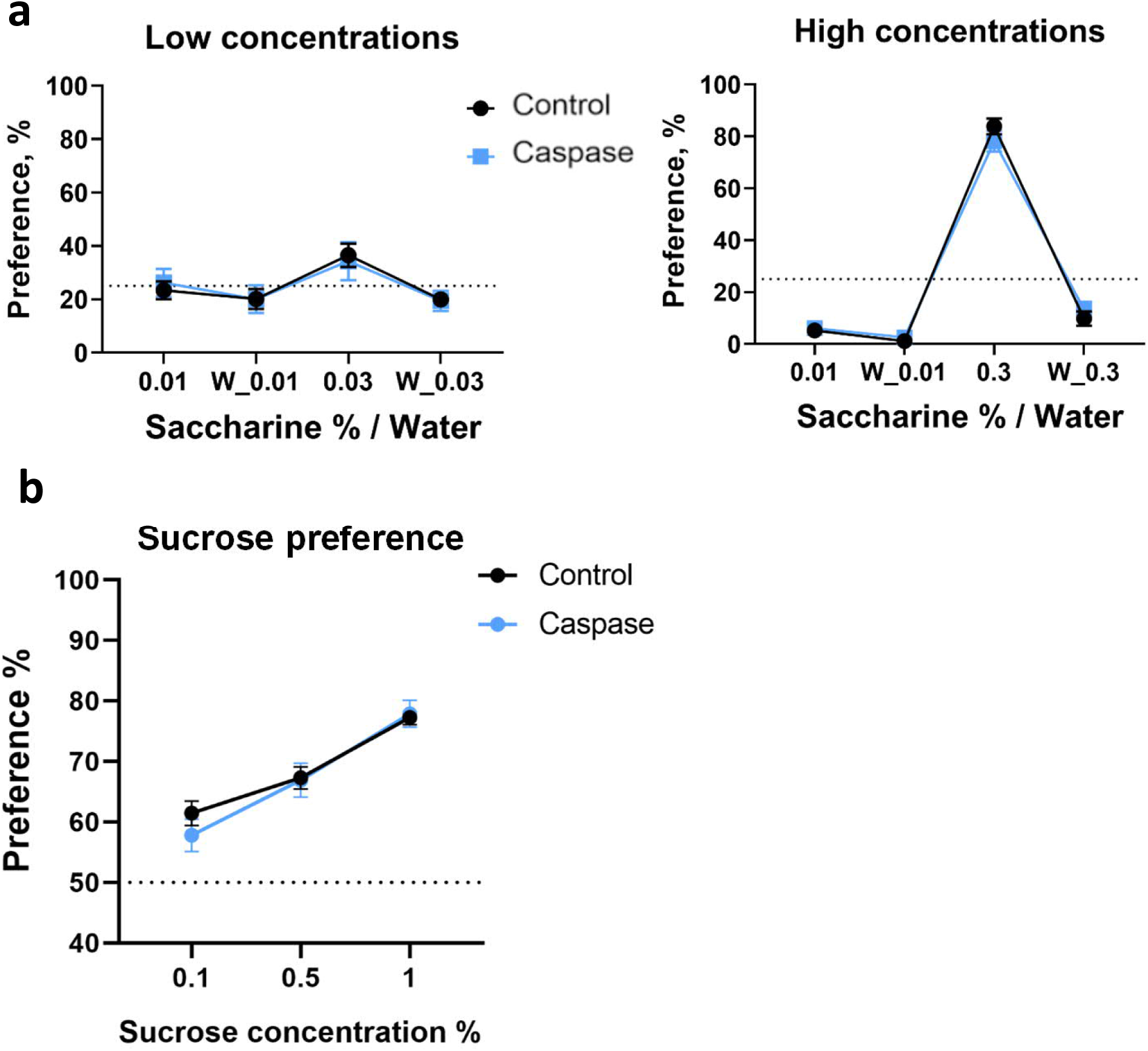
Deletion of the VTA Sst neurons did not affect natural reward preference or sensitivity. Graphs show preference in % (number of licks to a certain bottle/number of total licks, Y-axis) to different saccharine concentrations over water (X-axis). **a.** Preference to different saccharine concentrations or water in the corresponding corner in the Intellicage system did not reveal any significant differences between the treatment groups for low saccharine concentrations (F(1,24)=0.698, p=0.413) or to high ones (F(1,24)=0.045, p=0.834). The dashed line shows a 25% preference rate. **b**. Similarly, sucrose preference in the two-bottle choice test in individual cages did not reveal any differences (concentration x treatment: F(2,66) = 0.362, p = 0.688). The dashed line shows a 50% preference rate. Data are shown as mean ± SEM.

**Figure S11.**
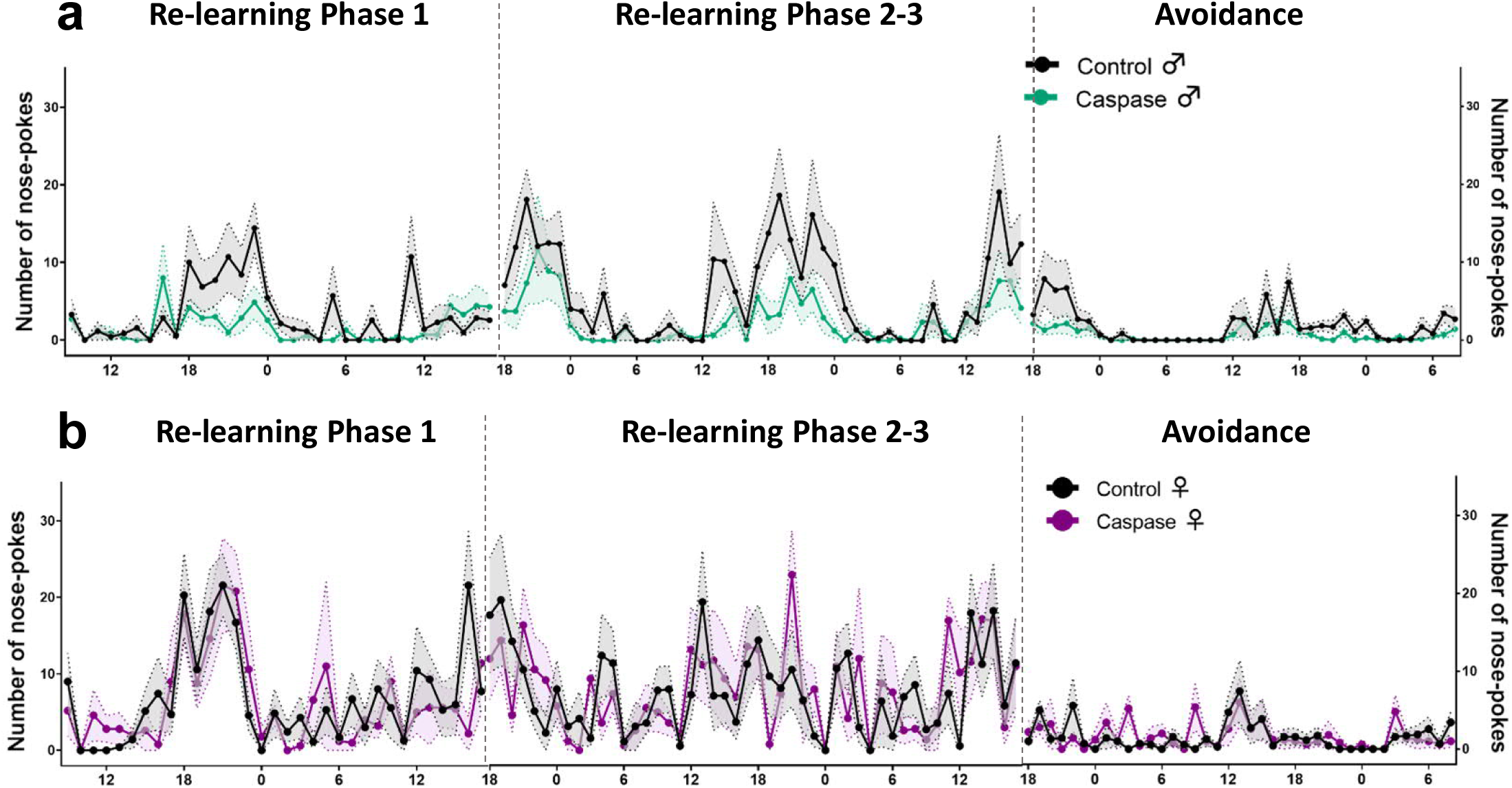
Re-learning new rules in control and caspase mice and introduction of air-puffs in the reward-related task. The X-axis shows the number of nose-pokes to the saccharine bottles per hour, Y-axis shows daily hours (lights on 6-18). **a.** Nose-poking dynamics in male mice. Although there was a clear tendency in VTA^Sst^- caspase male mice to be less active in nose-poking to the saccharine corner in all phases of the re-learning-avoidance test, no statistically significant differences were detected between the groups (see Table S2). **b.** Nose-poking dynamics in female mice showed no differences between the groups. Data are shown as mean ± SEM.

