## Supplementary Table 1 for "Somatostatin-expressing neurons in the ventral tegmental area innervate specific forebrain regions and are involved in the stress response"

### Intellicage

Supplementary Table 1

#### Home cage activity

*Number of nose-pokes* RM - repeated measures

| | | | | | | | | |
|---|---|---|---|---|---|---|---|---|
| Adaptation (14 hours) | RM-2 way ANOVA | F | p value | | post-hoc | | | |
| | Sex | F(1,22)=24.45 | <0.001 | **** | males | | females | |
| | Treatment | F(1,22)=4.543 | 0.044 | * | 0.698 | ns | 0.004 | *** |
| | Sex*Treatment | F(1,22)=7.085 | 0.014 | * | | | | |
| Day 2 (24 hours) | RM-2 way ANOVA | F | p value | | post-hoc | | | |
| | Sex | F(1,22)=5.751 | 0.025 | * | males | | females | |
| | Treatment | F(1,22)=2.770 | 0.11 | ns | 0.647 | ns | 0.03 | ** |
| | Sex*Treatment | F(1,22)=3.133 | 0.091 | ns | | | | |
| Day 3 (24 hours) | RM-2 way ANOVA | F | p value | | post-hoc | | | |
| | Sex | F(1,22)=18.315 | <0.001 | **** | males | | females | |
| | Treatment | F(1,22)=4.884 | 0.038 | * | 0.647 | ns | 0.002 | ** |
| | Sex*Treatment | F(1,22)=8.043 | 0.01 | * | | | | |
| All 3 days (62 hours) | RM-2 way ANOVA | F | p value | | post-hoc | | | |
| | Sex | F(1,22)=27.329 | <0.001 | **** | males | | females | |
| | Treatment | F(1,22)=7.409 | 0.012 | * | 0.695 | ns | <0.001 | **** |
| | Sex*Treatment | F(1,22)=10.618 | 0.004 | ** | | | | |

*Number of corner visits*

| | | | | | | | | |
|---|---|---|---|---|---|---|---|---|
| Adaptation (14 hours) | RM-2 way ANOVA | F | p value | | post-hoc | | | |
| | Sex | F(1,22)=4.53 | 0.045 | * | males | | females | |
| | Treatment | F(1,22)=2.131 | 0.158 | ns | 0.892 | ns | 0.047 | * |
| | Sex*Treatment | F(1,22)=2.707 | 0.114 | ns | | | | |
| Day 2 (24 hours) | RM-2 way ANOVA | F | p value | | post-hoc | | | |
| | Sex | F(1,22)=0.013 | 0.909 | ns | males | | females | |
| | Treatment | F(1,22)=0.069 | 0.795 | ns | 0.388 | ns | 0.258 | ns |
| | Sex*Treatment | F(1,22)=2.104 | 0.161 | ns | | | | |
| Day 3 (24 hours) | RM-2 way ANOVA | F | p value | | post-hoc | | | |
| | Sex | F(1,22)=13.796 | 0.001 | *** | males | | females | |
| | Treatment | F(1,22)=1.443 | 0.242 | ns | 0.798 | ns | 0.076 | ns |
| | Sex*Treatment | F(1,22)=2.406 | 0.135 | ns | | | | |
| All 3 days (62 hours) | RM-2 way ANOVA | F | p value | | post-hoc | | | |
| | Sex | F(1,22)=5.758 | 0.025 | * | males | | females | |
| | Treatment | F(1,22)=0.182 | 0.182 | ns | 0.574 | ns | 0.026 | * |
| | Sex*Treatment | F(1,22)=4.62 | 0.043 | * | | | | |

*Number of licks*

| | | | | | | | | |
|---|---|---|---|---|---|---|---|---|
| Adaptation (14 hours) | RM-2 way ANOVA | F | p value | | post-hoc | | | |
| | Sex | F(1,22)=22.97 | <0.001 | **** | males | | females | |
| | Treatment | F(1,22)=0.633 | 0.435 | ns | 0.605 | ns | 0.556 | ns |
| | Sex*Treatment | F(1,22)=0.008 | 0.931 | ns | | | | |
| Day 2 (24 hours) | RM-2 way ANOVA | F | p value | | post-hoc | | | |
| | Sex | F(1,22)=0.001 | 0.97 | ns | males | | females | |
| | Treatment | F(1,22)=0.709 | 0.409 | ns | 0.388 | ns | 0.258 | ns |
| | Sex*Treatment | F(1,22)=0.272 | 0.608 | ns | | | | |
| Day 3 (24 hours) | RM-2 way ANOVA | F | p value | | post-hoc | | | |
| | Sex | F(1,22)=1.716 | 0.204 | ns | males | | females | |
| | Treatment | F(1,22)=1.802 | 0.193 | ns | 0.578 | ns | 0.206 | ns |
| | Sex*Treatment | F(1,22)=0.339 | 0.567 | ns | | | | |
| All 3 days (62 hours) | RM-2 way ANOVA | F | p value | | post-hoc | | | |
| | Sex | F(1,22)=2.046 | 0.167 | ns | males | | females | |
| | Treatment | F(1,22)=1.390 | 0.251 | ns | 0.647 | ns | 0.253 | ns |
| | Sex*Treatment | F(1,22)=0.307 | 0.585 | ns | | | | |

Conclusion:
caspase animals overall nose poke more, but do not drink more (number of licks)
only caspase females significantly nosepoke and visit corners more than control females
no signinfican difference in males between the treatment groups

#### Delay discounting

| | | | | |
|---|---|---|---|---|
| *Number of licks to saccharine bottle* | RM-2 way ANOVA | F | p value | |
| | Sex | F(1,22)=5.889 | 0.0239 | * |
| | Treatment | F(1,22)=0.032 | 0.859 | ns |
| | Sex*Treatment | F(1,22)=0.836 | 0.37 | ns |
| | Sex*Delay | F(6,132)=17.09 | <0.001 | **** |

Conclusion: no significant difference between Treatment groups of both sexes
males are ready to wait saccharine less time than females

#### Saccharine learning and unlearning (number of nose pokes)

| | | | | | | | | |
|---|---|---|---|---|---|---|---|---|
| Adaptation | RM-2 way ANOVA | F | | | | | | |
| | Sex | F(1,22)=23.854 | <0.001 | **** | | | | |
| | Treatment | F(1,22)=1.217 | 0.282 | ns | | | | |
| | Sex*Treatment | F(1,22)=0.417 | 0.525 | ns | | | | |
| Basic activity | RM-2 way ANOVA | F | | | | | | |
| | Sex | F(1,22)=53.712 | <0.001 | **** | | | | |
| | Treatment | F(1,22)=1.345 | 0.259 | ns | | | | |
| | Sex*Treatment | F(1,22)=2.094 | 0.162 | ns | | | | |
| Unlearning | RM-2 way ANOVA | F | | | post-hoc | | | |
| "prediction error" | Sex | F(1,22)=33.575 | <0.001 | **** | males | | females | |
| | Treatment | F(1,22)=0.522 | 0.478 | ns | 0.3 | ns | 0.064 | ns |
| | Sex*Treatment | F(1,22)=4.635 | 0.043 | * | | | | |

Conclusion: females are more active than males in both treatment groups
Caspase females have a tendency to unlearn slower than the control female group but it is not significant

#### Saccharine re-learning and avoidance (number of nose pokes)

| | | | | | | | | |
|---|---|---|---|---|---|---|---|---|
| Adapatation | RM-2 way ANOVA | F | | | post-hoc | | | |
| ("re-learning") | Sex | F(1,22)=11.964 | <0.001 | **** | males | | females | |
| | Treatment | F(1,22)=1.547 | 0.227 | ns | 0.193 | ns | 0.659 | ns |
| | Sex*Treatment | F(1,22)=0.323 | 0.576 | ns | | | | |
| Basic activity | RM-2 way ANOVA | F | | | post-hoc | | | |
| | Sex | F(1,22)=8.298 | 0.009 | ** | males | | females | |
| | Treatment | F(1,22)=2.082 | 0.163 | ns | 0.034 | * | 0.917 | ns |
| | Sex*Treatment | F(1,22)=2.557 | 0.124 | ns | | | | |
| Avoidance | RM-2 way ANOVA | F | | | post-hoc | | | |
| "air-puffs" introduced | Sex | F(1,22)=0.617 | 0.441 | ns | males | | females | |
| | Treatment | F(1,22)=1.939 | 0.178 | ns | 0.03 | * | 0.816 | ns |
| | Sex*Treatment | F(1,22)=3.026 | 0.096 | ns | | | | |
| Full dynamics | RM-2 way ANOVA | F | | | post-hoc | | | |
| | Sex | F(1,22)=11.964 | 0.002 | ** | males | | females | |
| | Treatment | F(1,22)=2.069 | 0.164 | ns | 0.047 | * | 0.982 | ns |
| | Sex*Treatment | F(1,22)=1.975 | 0.174 | ns | | | | |

Conclusion: females are more active than males in both treatment groups, but not after the air-puffs were introduced
No differences in the re-learning or avoidance rates between treatment groups.
