## Supplementary Table 2 for "Somatostatin-expressing neurons in the ventral tegmental area innervate specific forebrain regions and are involved in the stress response"

### FEAR CONDITIONING

Supplementary Table 2

#### FEAR ACQUISITION (conditioning)

| | | | | | |
|---|---|---|---|---|---|
| pcnt freezing (PF) | RM-2 way ANOVA | F | p value | post-hoc | post-hoc |
| | Sex | F(1,19)=0.14 | 0.906 ns | males | females |
| | Treatment | F(1,19)=0.499 | 0.488 ns | 0.071 ns | 0.017 * |
| | Sex*Treatment | F(1,19)=10.459 | 0.004 ** | tendency that | caspase freeze more |
| | Point*Sex*Treatment | F(6,114)=4.046 | 0.012 * | caspase freeze less | |
| freezing episodes (FE) | RM-2 way ANOVA | F | p value | | |
| | Sex | F(1,19)=0.951 | 0.342 ns | | |
| | Treatment | F(1,19)=0.419 | 0.525 ns | *no within Sexes* | |
| | Sex*Treatment | F(1,19)=5.159 | 0.035 * | | |
| PF aqusition only BREAKS | RM-2 way ANOVA | F | | post-hoc | post-hoc |
| pcnt freezing | Sex | F(1,19)=0.028 | 0.869 ns | males | females |
| | Treatment | F(1,19)=0.033 | 0.857 ns | 0.036 * | 0.038 * |
| | Sex*Treatment | F(1,19)=9.971 | 0.005 ** | caspase freeze less | caspase freeze more |
| FE aqusition only BREAKS | RM-2 way ANOVA | F | p value | | |
| | Sex | F(1,19)=1.368 | 0.257 ns | | |
| | Treatment | F(1,19)=0.120 | 0.733 ns | *no within Sexes* | |
| | Sex*Treatment | F(1,19)=4.828 | 0.041 * | | |

RM - repeated measures

*No difference in only Cue Sounds (CS) periods*

| | | | |
|---|---|---|---|
| | RM-2 way ANOVA | F | p value |
| pcnt freezing | Sex | F(1,19)=0.019 | 0.891 ns |
| | Treatment | F(1,19)=2.182 | 0.156 ns |
| | Sex*Treatment | F(1,19)=2.390 | 0.139 ns |

#### CONTEXT RETRIEVAL 9 DAYS AFTER

| | | | | | |
|---|---|---|---|---|---|
| pcnt freezing (PF) | 2 way ANOVA | F | p value | post-hoc | post-hoc |
| | Sex | F(1,21)=1.191 | 0.288 ns | males | females |
| | Treatment | F(1,21)=0.18 | 0.894 ns | 0.08 ns | 0.086 ns |
| | Sex*Treatment | F(1,21)=6.602 | 0.018 * | | |
| Freezing episodes (FE) | 2 way ANOVA | F | p value | post-hoc | post-hoc |
| | Sex | F(1,21)=0.25 | 0.876 ns | males | females |
| | Treatment | F(1,21)=3.269 | 0.085 ns | 0.622 ns | 0.01 * |
| | Sex*Treatment | F(1,21)=6.102 | 0.022 * | | caspase freeze more often |

#### CUE-INDUCED RETRIVAL AND EXTINCTION

| | | |
|---|---|---|
| RM-2 way ANOVA | F | p value |
| Sex | F(1,21)=4.156 | 0.54 ns |
| Treatment | F(1,21)=0.322 | 0.577 ns |
| Sex*Treatment | F(1,21)=1.823 | 0.191 ns |
| Cue*Treatment | F(20,420)=0.78 | 0.642 ns |
| Cue*Sex*Treatment | F(20,420)=1.040 | 0.41 ns |

*no difference in CS retrieval*
