## Supplementary Table 3 for "Somatostatin-expressing neurons in the ventral tegmental area innervate specific forebrain regions and are involved in the stress response"

### OTHER BEHAVIOURAL TESTS

Supplementary Table 3

| NOVELTY-INDUCED LOCOMOTION (60 min) | | | | |
|---|---|---|---|---|
| Distanced moved | RM-2 way ANOVA | F | p-value | |
| 10 min bins | Treatment | F(1,34) = 0.265 | 0.61 | ns |
| | Treatment*sex | F(1,34) = 0.203 | 0.655 | ns |
| | Treatment*time | F(5,170) =0.43 | 0.782 | ns |
| *No difference* | | | | |

RM - repeated measures

| LIGHT-DARK BOX TEST | | | | |
|---|---|---|---|---|
| Time spent in light (%) | 2 way ANOVA | F | p-value | |
| | Sex | F(1,34) = 3.957 | 0.055 | ns |
| | Treatment | F(1,34) = 1.75 | 0.195 | ns |
| | Treatment*sex | F(1,34) = 0.704 | 0.407 | ns |
| *No difference* | | | | |

| ELEVATED PLUS MAZE | | | | |
|---|---|---|---|---|
| Time spent in open arm (%) | RM-2 way ANOVA | F | p-value | |
| | Sex | F(1,34) = 2.656 | 0.112 | ns |
| | Treatment | F(1,34) = 0.072 | 0.79 | ns |
| | Treatment*sex | F(1,34) = 0.394 | 0.534 | ns |
| *No difference* | | | | |

| RUNNING WHEEL (RW) ACTIVITY | | | | |
|---|---|---|---|---|
| 3 Days activity | RM-2 way ANOVA | F | p-value | |
| | Sex | F(1,24) = 1.470 | 0.237 | ns |
| | Treatment | F(1,24) = 0.202 | 0.657 | ns |
| | Treatment*sex | F(1,24) = 1.574 | 0.222 | ns |
| | Treatment*time | F(67,1608) = 1.230 | 0.278 | ns |
| | Treatment*time*sex | F(67,1608) = 0.729 | 0.683 | ns |
| *No difference* | | | | |

| NOVELTY-SUPPRESSED FEEDING | | | | |
|---|---|---|---|---|
| Latency to eating | 2 way ANOVA | F | p-value | |
| | Sex | F(1,34) = 0.543 | 0.47 | ns |
| | Treatment | F(1,34) = 0.029 | 0.865 | ns |
| | Treatment * sex | F(1,34) = 3.862 | 0.058 | ns |
| *No difference* | | | | |

| FORCED SWIM TEST | | | | |
|---|---|---|---|---|
| Latency to immobilization | 2 way ANOVA | F | p-value | |
| | Sex | F(1,34) = 0.33 | 0.858 | ns |
| | Treatment | F(1,34) = 5.08 | 0.031 | * |
| | Treatment * sex | F(1,34) = 0.989 | 0.327 | ns |
| *Caspase display longer latency to first immobility* | | | | |
| Duration of immobiliation | RM-2 way ANOVA | F | p-value | |
| 2 min bins | Sex | F(1,34) = 0.611 | 0.611 | ns |
| | Treatment | F(1,34) = 0.263 | 0.214 | ns |
| | Treatment * sex | F(1,34) = 0.08 | 0.778 | ns |
| *No difference* | | | | |

Supplementary Table 3

| MORPHINE-INDUCED MOTOR SENSITIZATION | | | | |
|---|---|---|---|---|
| Induction | RM-2 way ANOVA | F | p-value | |
| Distance moved | Sex | F(1,32) = 0.04 | 0.95 | ns |
| 10 min bins | Treatment | F(1,32) = 0.034 | 0.854 | ns |
| | Treatment * sex | F(1,32) = 0.12 | 0.731 | ns |
| | Treatment * time | F(23,736) = 0.153 | 0.917 | ns |
| | Treatment * time * sex | F(23,736) = 0.146 | 0.922 | ns |
| *No difference* | | | | |
| Challenge | RM-2 way ANOVA | F | p-value | |
| **Δ** Distance moved Challenge-Induction | Sex | F(1,32) = 3.0 | 0.93 | ns |
| 10 min bins | Treatment | F(1,32) = 12.014 | 0.002 | ** |
| | Treatment * sex | F(1,32) = 0.654 | 0.425 | ns |
| | Treatment * time | F(23,736) = 1.649 | 0.024 | * |
| | Treatment * time * sex | F(23,736) = 2.915 | 0.166 | ns |
| *Upregulation of morphine motor sensitization* | | | | |

| SUCROSE PREFERENCE | | | | |
|---|---|---|---|---|
| Sucrose preference (%) | RM-2 way ANOVA | F | p-value | |
| | Sex | F(1,33) = 0.413 | 0.413 | ns |
| | Treatment | F(1,33) = 0.0 | 0.995 | ns |
| | Treatment * sex | F(1,33) = 2.347 | 0.135 | ns |
| | Treatment * concentration | F(2,66) = 0.362 | 0.688 | ns |
| | Treatment * concentration * sex | F(2,66) = 2.317 | 0.109 | ns |
| *No difference* | | | | |
